# *SAS_MoCa*: a software for small-angle scattering data analysis of large unilamellar vesicles

**DOI:** 10.64898/2026.06.29.735169

**Authors:** Enrico F. Semeraro, Georg Pabst

**Affiliations:** University of Graz, Institute of Molecular Biosciences, NAWI Graz, 8010 Graz, Austria; Field of Excellence BioHealth – University of Graz, Graz, Austria; BioTechMed Graz, 8010 Graz, Austria

**Keywords:** Small-angle X-ray scattering, Large unilamellar vesicles, Bayesian inference, Membrane structure, Open-source software

## Abstract

Small-angle X-ray or neutron scattering (SAXS/SANS) analysis of large unilamellar vesicles (LUVs) is often limited by high-dimensional bilayer models and the lack of dedicated, statistically rigorous workflows. Here, we introduce *SAS_MoCa*, an open-source Python package that integrates a compositional scattering density profile (SDP) description of lipid bilayers with a separated form factor (SFF) treatment of vesicle size and polydispersity, and couples these highly parameterized models to an adaptive thermodynamic simulated annealing algorithm formulated within a constrained Bayesian framework. *SAS_MoCa* enables users to incorporate quantitative prior information from, e.g., previous SAXS/SANS studies, dynamic light scattering, NMR, or molecular simulations, and returns full posterior parameter distributions, uncertainties (reported as medians and median absolute deviations) and correlations even from single SAXS curves. Validation on POPC, POPE and DMPC SAXS-only data demonstrates that the method yields reproducible structural parameters with uncertainties comparable to joint SAXS/contrast-variation SANS analyses. The modular architecture of *SAS_MoCa* facilitates extension to additional lipid systems and future joint SAXS/SANS or SANS-only applications.

**Synopsis:** *SAS_MoCa* is an open-source Python package for quantitative analysis of small-angle scattering (SAS) data from large unilamellar vesicles, combining a compositional scattering density profile model of lipid bilayers with a separated form factor description of vesicle size and polydispersity in a Bayesian framework. It yields full posterior parameter distributions, uncertainties and correlations from single SAS curves, enabling reproducible and extensible high-throughput membrane structural characterization.

## 1 Introduction

Lipid vesicles of defined size and composition are highly popular platforms in fundamental life science/soft matter research and numerous biotech applications (Pabst et al., 2014). In particular organelle-sized vesicles, so called large unilamellar vesicles (LUVs) with sizes around 100 nm, often serve as minimal models of cellular membranes, extracellular vesicles, or lipid nanoparticles for diverse drug-delivery formulations (see e.g. (Chacko et al., 2020)). Small-angle scattering (SAS) stands as one of the most powerful and non-invasive techniques for probing LUV structure at mesoscopic length scales (Semeraro et al., 2024). However, extracting meaningful structural parameters from SAS data typically demands significant expertise in scattering theory, model selection, and statistical validation.

Currently, popular SAS analysis tools such as *SasView* (Krzywon et al., 2025) lack a bespoke workflow tailored specifically to LUVs, while a previously dedicated online tool, *Vesicle Viewer* (Lewis-Laurent et al., 2021), is no longer maintained. Alternatively, the scattering length density profile of lipid bilayers can be analyzed via an heuristic combination of Gaussian curves with the routine *LIPMIX*, embedded in the ATSAS software (Konarev et al., 2021), extending a previously reported full *q*-range analysis of multilamellar and unilamellar vesicles (Pabst et al., 2000; Pabst et al., 2003). However, none of these tools combine compositional information of the lipid bilayer with a vesicle size and polydispersity analysis. Consequently, high fidelity scattering data analysis of LUVs presents a pressing bottleneck and significant barrier for progress in fundamental biological membrane research and biotechnological applications of these systems.

To address these challenges, we present *SAS_MoCa*, an open-source tool enabling Bayesian-constrained compositional analysis of LUV small-angle X-ray scattering (SAXS) data (Semeraro & Pabst, 2026). Specifically, *SAS_MoCa* provides a multiscale analysis that integrates the scattering density profile (SDP) model for high-resolution bilayer structure (Kučerka et al., 2008) and the separated form factor (SFF) model for vesicle size and size polydispersity (Pencer et al., 2006). The package provides a unified, open-source framework that streamlines data import, curve fitting, result aggregation, and visualization, thereby lowering the barrier to entry for non-specialists in scattering analysis.

*SAS_MoCa* enables rigorous and efficient statistical analysis of small-angle scattering data. At the core of the package is an adaptive stochastic minimization engine tailored to explore the high-dimensional parameter space characteristic of SDP modeling of LUVs. This engine is formulated within a Bayesian framework, allowing users to incorporate prior information—such as bilayer thickness or vesicle radius—and to obtain full posterior distributions for all adjustable parameters. While deterministic optimizers (e.g., Levenberg–Marquardt) offer rapid convergence to local minima, they lack the capacity to quantify parameter uncertainties or explore complex likelihood landscapes. *SAS_MoCa* bridges this gap by implementing constrained Bayesian inference that provides full statistical characterization—median values, credible intervals, and parameter correlations—in a matter of minutes even on a standard laptop.

*SAS_MoCa* thus offers a unified framework for *integrating SAXS measurements with diverse quantitative priors*. Instead of relying exclusively on the scattering curve, users can combine it with: (i) literature values from previous joint SAXS/SANS studies of analogous lipid systems; (ii) independent experimental measurements such as, e.g., Dynamic Light Scattering (DLS) for vesicle size; and (iii) computational results providing high-resolution structural information beyond the intrinsic resolution of SAS. This enables knowledge transfer from well-characterized systems to new experiments, and establishes a general, rigorous route for incorporating external constraints into scattering-based structural analysis.

Finally, the software architecture is deliberately modular, making it straightforward to extend the package to alternative scattering models and to tackle small-angle neutron scattering (SANS) data, including contrast-variation and joint SAXS/SANS analyses.

This manuscript is organized as follows. First, we present the scattering models and the Bayesian-stochastic fitting engine implemented in *SAS_MoCa*. We then validate the methodology using experimentally measured scattering data from LUVs of varying compositions, and assess its performance and limitations. Finally, we summarize the main conclusions, and outline avenues for future developments and extensions.

## 2 Program Overview

### 2.1 Scattering Form Factor

The *SAS_MoCa* repository currently includes scattering form factors designed to analyze the bilayer structure and size of LUVs. We assume that the vesicles are fully hydrated within the lamellar fluid phase, L_*α*_, and freely float in pure water. The scattering intensity of a suspension of LUVs is expressed as:

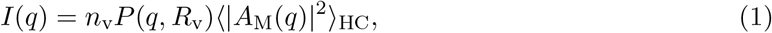

where *n*_v_ is the number density of LUVs; *P* (*q, R*_v_) represents the SFF contribution of the spherical LUV frame of radius *R*_v_, including polydispersity effects (Pencer et al., 2006); and the scattering amplitude *A*_M_(*q*) describes the one-dimensional lipid bilayer contribution according to the SDP formalism (Kučerka et al., 2008; Frewein et al., 2021). The averaging operator ⟨· · · ⟩_HC_ denotes integration over a normal distribution of hydrocarbon layer thicknesses (Frewein et al., 2021). Detailed derivations of the model implementation are provided in Appendix A.

While the SFF model considers LUVs as rigid spherical frames with a given size distribution, the SDP approach treats each lipid as a collection of quasi-molecular groups (see Fig. 1), with each group contributing individually to the scattering signal (Kučerka et al., 2008). As the SDP is a *compositional* model, the specifics of *A*_*M*_ (*q*) depend directly on lipid composition (see Fig. 1 and Appendix A). *SAS_MoCa* currently implements the compositional design for the following lipid systems: POPC (1-palmitoyl-2-oleoyl-*sn*-glycero-3-phosphocholine), DMPC (1,2-dimyristoyl-*sn*glycero-3-phosphocholine), or POPE (1-palmitoyl-2-oleoyl-*sn*-glycero-3-phosphoethanolamine).

**Figure 1.**
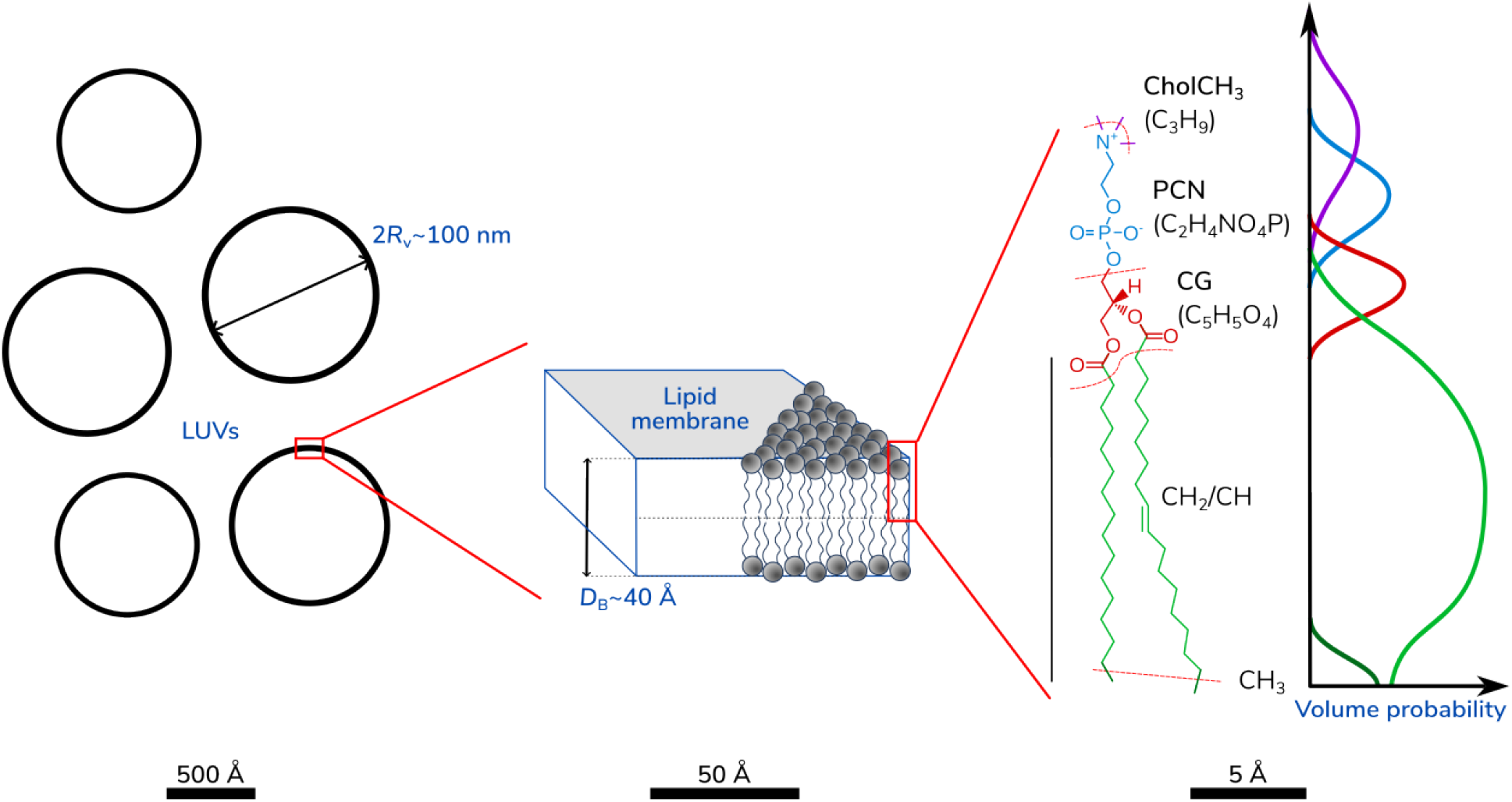
Schematic representation of the multi-scale SFF-SDP model, from LUVs’ size to the parsing of quasi-molecular groups in, e.g., POPC lipids (Kučerka et al., 2011).

#### 2.1.1 Numerical Implementation

While Eq. 1 defines the theoretical scattering intensity, *SAS_MoCa* implements this calculation within the `LUV_POPC` class (and analogous classes for DMPC and POPE) through a sequential modular workflow of component definitions. For users navigating the source code (e.g., `sasmoca/models/LUV_POPC.py`), the implementation logic follows these key steps:

1. **Initialization**: The constructor parses input parameters, *y*, computes lipid-specific volumes (e.g., *V*_L_, *V*_HC_) and loads the scattering lengths, *b*, pertinent to the quasi-molecular groups of the specific compositional design from `sasmoca/models/constants.py` (see Tab. S1). Volumes and *b*-values are combined in scattering length density, Δ*ρ*, values as functions of temperature and composition. The initialization further establishes the LUV geometric framework, including vesicle polydispersity and hydrocarbon layer thickness distributions.
2. **Amplitude Calculation (**`Am`): The core membrane scattering amplitude *A*_M_(*q*) is assembled by summing contributions from quasi-molecular groups. The code distinguishes between smooth rectangular profiles (acyl chains, hydration layers) computed via error-function Fourier transforms (`FTreal_erf`) and Gaussian profiles (head-groups, terminal methyls) using `FTreal_gauss` (see Appendix A). An integration grid over the hydrocarbon thickness distribution is generated to account for thickness fluctuations ⟨`|`*A*_M_(*q*)`|`^2^⟩_HC_.
3. **Intensity Aggregation**: The scattered intensity is computed according to Eq. 1. The method `intensity()` then returns the total model curve *I* = *NI*(*q*) + *C*(*q*), where *N* is a normalization scaling factor. Notably, the background term *C*(*q*) = *Const*. × {*v/*[1 + exp(−*κ*(*q* − *q*_0_))] + (1 − *v*)} incorporates a sigmoidal transition at high-*q* values. This functional form accounts for the typical unidentified scattering contributions at high-*q* values without obscuring the structural signal in the mid-*q* range where the form factor oscillations reside (Semeraro et al., 2024). Current settings are optimized for *q*_0_ = 0.1 Å^−1^, *κ* = 8 Å and *v* = 0.99.

Note that the lipid volume, *V*_L_, and scattering length density of suspension water, Δ*ρ*_sol_, are automatically calculated for the sample temperature, *T*, by interpolating reported data. For the PC, PE and PG species considered here, lipid volumes are obtained by linear interpolation in the range 20−60 ^°^C using the data reported in (Kučerka et al., 2011), (Kučerka et al., 2015) and (Pan et al., 2012), respectively. The scattering length density of the suspending water is instead calculated through the water molecule volume, which in turn is obtained via a polynomial interpolation in the range 1−100 ^°^C using standard tables (see, e.g., (Harvey, 1998)). Linear coefficients for lipid volumes and polynomial coefficients for water volumes are listed in `sasmoca/models/constants.py`.

### 2.2 Minimization Algorithm

To optimize the large number of adjustable parameters of SDP-based LUV model, we employ a Monte Carlo approach that combines efficient *χ*^2^ minimization with rapid convergence. The core of our algorithm consists of a series of Thermodynamic Simulated Annealing (TSA) minimization runs, adapted from (de Vicente et al., 2003). Each iteration is independent, allowing for parallel execution. The workflow of the program is depicted in Figure 2.

**Figure 2.**
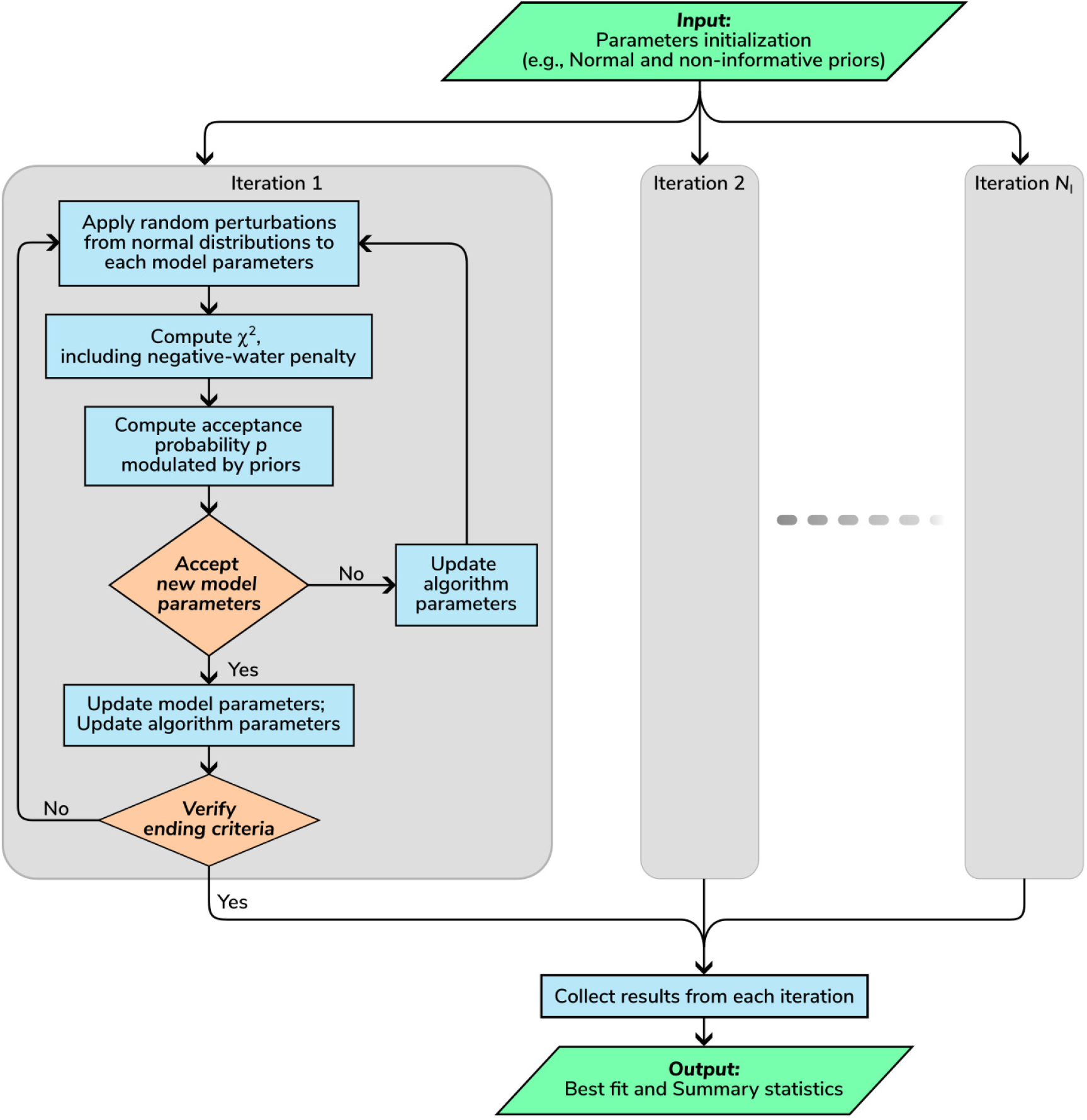
Flowchart of the *SAS_MoCa* minimization algorithm. The process begins with parameter initialization, proceeds through iterative TSA loops involving perturbations and acceptance checks, and concludes with the collection of best-fit parameters and statistical analysis.

#### 2.1.1 Initialization

In *SAS_MoCa*, we initialize each adjustable parameter, *y*, based on an initial guess and prior knowledge. Users can define two types of priors:

- **Non-informative priors, i.e**., **hard boundaries**: Users can set minimum (*y*_min_) and maximum (*y*_max_) limits. This corresponds to a rectangular prior distribution, referred to as a *non-informative* prior, where all values within the range are equally probable.
- **Informative priors**: Users can specify a normally distributed prior, *π*(*µ*_*y*_,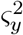), by setting its mean, *µ*_*y*_, and relative standard deviation, *ς*_*y*_*/µ*_*y*_. By default we impose additional hard boundaries at ±5*ς*_*y*_ around *µ*_*y*_ for computational stability.

Beyond model parameters, users can configure global options, such as the total number of TSA iterations, *N*_*I*_. These iterations can be executed sequentially or distributed across multiple parallel processes.

#### 2.2.2 Single TSA Iteration

Each TSA iteration begins by calculating the initial reduced 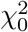 from the starting parameter set. The algorithm then enters an iterative loop wherein adjustable parameters are perturbed, an acceptance probability is computed, and the new parameter set is evaluated. This loop continues until specific termination criteria are met.

##### Apply random perturbations

In each step, the current model parameters are modified by small random perturbations. Specifically, every adjustable parameter *y*_*i*_ is shifted by a value drawn from a normal distribution centered at zero, with a standard deviation equal to *β*(*y*_*i*,max_ − *y*_*i*,min_)*/*6, where *β* is a tunable scale factor that adapts dynamically throughout the minimization.

##### Negative water penalty

We incorporate a penalty term into the reduced *χ*^2^ calculation to suppress configurations that yield non-physical water volume fractions within the lipid bilayer. In the SDP formalism, the headgroup hydration layer volume is obtained by subtracting the volumes of the quasi-molecular lipid groups located in the headgroup region from the total hydration shell volume. If the fitted parameters imply a lipid headgroup volume larger than the available shell volume, the residual water volume becomes negative and thus unphysical. To detect and penalize such cases systematically, we implement the following procedure.

First, the `negative_water()` method discretizes the bilayer profile and counts the number of grid points, *N*_neg_, where the local water volume fraction falls below a threshold of −0.01. This count is then converted into a multiplicative penalty factor that is applied to the reduced *χ*^2^ prior to the acceptance step. The effective 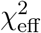 is defined as

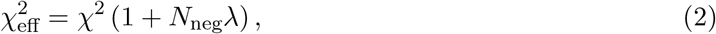

where *λ* is a user-defined scaling factor (default `neg_water_scale: 5`, set in the YAML configuration file `settings.yml`). Because the acceptance probability depends exponentially on 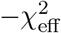 (Eq. 3), even moderate values of *λ* strongly disfavor parameter sets with non-physical water distributions, guiding the TSA algorithm away from such regions without introducing hard constraints that could trap the optimization.

##### Compute probability of acceptance

We evaluate each trial set of model parameters based on the effective reduced 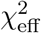 and the distances from the prior centers. For the *k*-th iteration within a TSA loop, the acceptance probability *p*_*k*_ is given by:

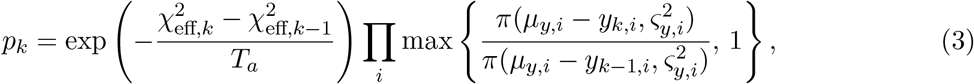

where *T*_*a*_ is the annealing temperature parameter, and *π*(*µ*_*y,i*_, 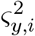) represents the normal prior for the *i*-th adjustable model parameter. The first factor corresponds to the standard thermodynamic simulated annealing criterion (de Vicente et al., 2003), while the product over *i* implements the prior-based acceptance modification described in (Frewein et al., 2019).

##### Evaluate new set of parameters

We accept or reject the proposed set of randomly perturbed parameters by comparing a random value *p*_0_ ∈ [0, 1] against *p*_*k*_:

- If 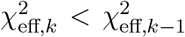: the *k*-th set is accepted. It replaces the previous parameter set and is stored as the best set of parameters. Furthermore, if 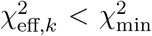 (the lowest *χ*^2^ found so far), we update 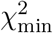.
- If 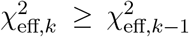 and *p*_0_ *< p*_*k*_: the *k*-th set is accepted and replaces the previous one. However, the best parameters and 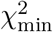 remain unchanged.
- If 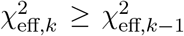 and *p*_0_ ≥ *p*_*k*_: the *k*-th set is rejected, and the previous set remains unchanged.

We also use this assessment to update algorithm parameters. Specifically, we compute the acceptance/rejection rates to adjust the TSA temperature *T*_*a*_ (de Vicente et al., 2003), and we dynamically modify the scale factor *β* to maintain an acceptance rate between 60% and 70%.

##### Check for ending criteria

The TSA loop terminates when a user-defined maximum number of repetitions is reached (default `maxcount: 100000`, configured in `settings.yml`) or convergence is detected. Convergence is determined by monitoring the cumulative likelihood of approaching a user-defined target 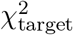. Specifically, we check if:

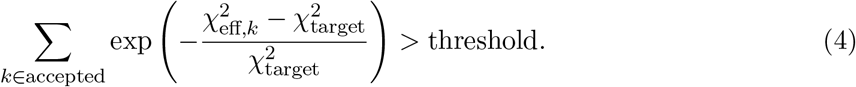

where the threshold (default `conv_threshold: 4`) can be configured in `settings.yml`. Note that if 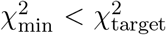, we automatically update 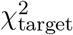 to the new minimum, ensuring the convergence criterion adapts to the best solution found.

#### 2.2.3 Collecting Best Results and Summary Statistics

Finally, we collect the final, best set of parameters and 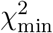 from each independent TSA run to compute summary statistics. This collection yields a posterior-like distribution for each model parameter, from which we calculate median values, the Median Absolute Deviation (MAD) from medians, and the Pearson correlation coefficients.

### 2.3 Statistical Background: Bayesian Inference and Information Update

The core philosophy of *SAS_MoCa* rests on the Bayesian principle of *updating beliefs*. In traditional least-squares fitting, the goal is often to find a single set of parameters that minimizes the discrepancy between model and data, implicitly assuming the data contains sufficient information to determine every parameter uniquely. However, in complex scattering models such as the SDP model, the scattering signal often lacks the resolution to constrain all physical parameters simultaneously.

Bayesian inference addresses this limitation by treating parameters as stochastic variables characterized by probability distributions. The analysis exploits the the Bayes’ theorem:

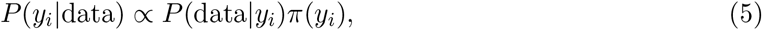

where *P* (*y*_*i*_|data) is the *posterior* distribution of the parameters *y*_*i*_ given the data, *P* (data|*y*_*i*_) is the *likelihood* (how well the model fits the data), and *π*(*y*) is the *prior* distribution representing our knowledge before ‘seeing’ the data (Ramachandran & Tsokos, 2015).

In the context of *SAS_MoCa*, the stochastic minimization algorithm samples this posterior distribution. The result of this sampling process reveals the extent to which the experimental data has *updated* our initial assumptions. We can distinguish two limiting behaviors based on the degree of this update:

1. **Data-Dominated Updates (‘Sensitive” Parameters):** When the experimental signal is strong and informative regarding a specific parameter *y*_*i*_, the likelihood term *P* (data|*y*_*i*_) is a narrow and bell-shaped distribution. The posterior distribution *P* (*y*_*i*_|data) is dominated by the data, regardless of the shape of the prior. Even if we shift the prior mean significantly, the resulting posterior remains stable and centered near the value dictated by the scattering curve. In this regime, the data has successfully “updated” the prior, extracting new information that was not present in our initial guess.
2. **Prior-Dominated Updates (“Non-Sensitive” Parameters):** Conversely, when the data contains insufficient information to constrain a parameter (e.g., due to limited *q*-range, low contrast or high noise), the likelihood distribution is rather featureless over a wide range of possible parameter values. In such cases, the posterior distribution closely mirrors the prior distribution, that is *P* (*y*_*i*_|data) ≈ *π*(*y*_*i*_). The Bayesian update fails to shift the distribution because the analysis cannot discriminate between different values. Here, the final result is essentially a confirmation of our prior assumption, not a discovery derived from the experiment.

This framework changes the interpretation of the output. The primary goal of *SAS_MoCa* is not merely to provide point estimates such as medians and deviations, but to visualize the *shift* between prior and posterior. By inspecting the posterior histograms alongside the priors (as demonstrated in section 4), users can immediately assess whether a parameter was truly determined by the experiment or constrained by external knowledge. If the posterior overlaps significantly with the prior, either the data confirm the previous belief or the user must acknowledge that the result relies on the validity of that prior. If the posterior is distinct from the prior, the experiment has provided independent structural insight. This distinction empowers a more rigorous workflow: users are encouraged to test the robustness of their results by varying the priors, either informative Gaussian or non-informative rectangular ones. If a parameter is truly sensitive, its posterior should remain stable under moderate prior shifts; if it is non-sensitive, the posterior will track the prior variations. This diagnostic capability is central to avoiding spurious conclusions in under-constrained systems.

## 3 Implementation / Program Description

*SAS_MoCa* is an open-source Python 3.12 software package distributed under the BSD-3-Clause license, available at https://github.com/PabstLab/SAS_MoCa. It runs via the command line on Linux, Windows, and macOS operating systems.

### 3.1 Architecture and Dependencies

The software architecture follows a modular, object-oriented design comprising three primary packages: `in_out` handles I/O routines, including data loading and result visualization; `moca` implements the TSA optimization engine; and `models` contains the scattering models for various LUV systems. The package currently includes SDP descriptions for POPC, DMPC, and POPE LUVs (see Eq. 1), which have been thoroughly tested. These are respectively implemented in the `LUV_POPC`, `LUV_DMPC` and `LUV_POPE` classes. This modular design ensures that the minimization algorithm operates independently from the model definitions, facilitating the extension to new membrane systems.

The core Python modules utilized are `pyyaml` for configuration input, `numpy` and `scipy` for scientific computing, `multiprocess` (version 0.70) for parallel computation, and `pandas` and `matplotlib` for data collection and visualization. No proprietary dependencies are required.

*SAS_MoCa* is designed as a portable Python package that runs without installation, ensuring consistent behavior across different environments. We recommend deploying the software within the dedicated `conda` environment managed via the provided `environment.yml` file to handle dependencies.

### 3.2 Input and Execution

Analysis is initiated via the command line interface (CLI):

`python <path-to-sasmoca.py>/sasmoca.py input-parameters.yml`

The input YAML file `input-parameters.yml` contains two blocks of information: `config` and `parameters`, which are needed to initialize and run the data analysis.

#### 3.2.1 Configuration Block (config)

The primary control settings are defined in the `config` block of the YAML input file (see Tab. 1). Crucially, this block specifies the target scattering or compositional model in terms of the SDP description (e.g., `LUV_POPC`), the experimental data path, and global minimization parameters; see section 4 for information how to handle lipid mixtures. Key parameters include `iterations` (*N*_*I*_), which determines the number of independent TSA runs to build statistical distributions, and `processes`, which enables parallel execution to leverage multi-core architectures. The `target-X2` parameter acts as a convergence guide (see Eq. 4).

#### 3.2.2 Parameter Initialization and Priors (parameters)

The core flexibility of *SAS_MoCa* lies in its `parameters` block, which allows for granular control over each adjustable variable using a compact list syntax:

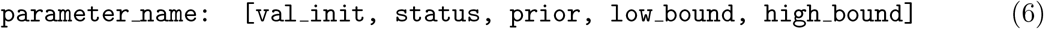

where:

- `val init`: The starting guess for the optimization routine or, if there is an associated Gaussian prior, the *µ*_*y*_ value.
- `status`: `‘on’` activates the parameter for fitting; `‘off’` fixes it.
- `prior`: The relative standard deviation *ς*_*y*_*/µ*_*y*_ for an informative Gaussian prior. If set to `Null`, only hard boundaries are applied (rectangular, non-informative prior).
- `low/high bound`: Hard boundaries for non-informative priors. If `prior` is active, it automatically overrides `low/high bound` values and sets additional bounds to *µ*_*y*_ `±` 5*ς*_*y*_. In this case, `low/high bound` can also be set to Null.

This syntax allows the user to specify both classical hard boundaries based on educated guesses and informative priors derived from literature (e.g., known head-group volumes) or complementary techniques (e.g., DLS-derived vesicle radii) directly within the fitting loop. For example, the line `R_v: [450, on, 0.10, Null, Null]` defines a Gaussian prior for the vesicle radius centered at the initial value 450 Å with a relative standard deviation of 10%. In contrast, `Z: [10, on, Null, 5, 20]` sets an initial value of 10 and uses a non-informative (uniform) prior between 5 and 20.

Complete templates are provided in the repository’s `examples` folder, demonstrating how to initialize parameters and priors for POPC, POPE and DMPC LUV systems.

### 3.3 Output

The output consists of a collection of lightweight ASCII files and PNG plots (totaling ~ 1 MB). Curve fitting results are summarized in two categories:

- **Visualization outputs**: `plot.png` (curve fitting overlay); `plot_SDP-SLD.png` (volume distribution of lipid quasi-molecular groups and the associated scattering length density profile); `plot_histograms.png` (sampling distributions of adjustable parameters); `plot_histogram X2.png` (sampling distribution of 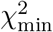 values); and `plot_correlations.png` (Pearson correlation coefficient matrix for adjustable parameters).
- **Fitting results and statistical summaries**: `results recap.dat` (summary statistics over the collection of iterations); `iterations collection.dat` (collection of best parameters from each iteration); `plot_intensity.dat` (ASCII file for custom re-plotting of the fitted curve); `plot_SDP-SLD.dat` (ASCII file for custom re-plotting of the volume distribution and/or SLD profile); and `correlations.dat` (Pearson correlation coefficient matrix).

Output files are automatically saved in a newly created result folder. Each ASCII output file includes embedded metadata documenting the configuration used, ensuring full reproducibility. These files are formatted for easy reading and can serve as input for subsequent analysis workflows.

## 4 Validation and Benchmarking

To validate *SAS_MoCa*, we applied the software to structural analyses of LUVs composed of three distinct phospholipid systems: POPC, DMPC, and POPE. The validation strategy combined benchmarking against reference POPC data taken from (Frewein et al., 2021), with fresh analysis of newly acquired SAXS data for water suspensions of LUVs consisting of either DMPC or POPE.

SAXS POPE and DMPC data were recorded at BM29, ESRF, Grenoble, France (proposal No. MX-2282). Samples were measured at *T* = 50 ^°^C in a flow-through quartz capillary of a 1 mm path length. Data reduction and normalization were performed by the automated pipeline system (Tully et al., 2023). All LUV samples (including the reference POPC data) were prepared with an extrusion diameter of 100 nm following standard protocols detailed in (Frewein et al., 2021). Note that all samples include a small percentage (5 − 10 mol%) of PG (phosphoglycerol) lipids with matching chain groups. The doping with PGs aids the fabrication of 100 nm-sized LUVs by extrusion through polycarbonate membranes (Scott et al., 2019). In *SAS_MoCa*, the POPC and POPE compositional models account for, respectively 5 and 10 mol% POPG, while DMPC contains 5 mol% DMPG. These small PG lipid percentages are included in the SDP model by using a weighted average of the quasi-molecular group volumes and scattering lengths. Details about the parsing of quasi-molecular groups for different species including PG lipids are in section S1 of the SI and in (Kučerka et al., 2011; Kučerka et al., 2015; Pan et al., 2012).

### 4.1 Validation of LUV Compositional Models

For the analysis of reference POPC SAXS data we initialized the curve fitting with a number of Gaussian priors selected from previously reported analyses (Kučerka et al., 2011; Frewein et al., 2021). Compared to those studies, we increased the degrees of freedom by expanding the number of adjustable parameters, leveraging dynamic light scattering (DLS) measurements for LUV radius determination, and estimates for lipid head-group volumes (Nagle et al., 2019). When previous measurements were not available, we used educated guesses to set non-informative priors via the hard-boundaries. The number of iterations was set to *N*_*I*_ = 300 to ensure reproducibility (section 4.3). Table 2 summarizes the parameters of our scattering model, their descriptions, and the prior initialization values. Note that the errors reported in (Kučerka et al., 2011) are fixed to 2% for all parameters. For the adjustable parameters *r*_PCN_, *r*_CG_, *r*_12_, and *r*_32_ we opted to increase the uncertainty to 5% to further challenge the validation of *SAS_MoCa*. We applied the same strategy to the analysis of POPE and DMPC data. When possible, we initialized Gaussian priors from (Kučerka et al., 2015; Frewein et al., 2023) for POPE and (Kučerka et al., 2011) for DMPC. Tables S2 and S4 summarizes the model parameters, their descriptions, and the prior initialization values.

**Table 1.** Summary of configuration options in the config block.

| Option | Description |
| --- | --- |
| `datafile` | Path to ASCII file (3 columns: $q_{\text{exp}}$, $I_{\text{exp}}$, $\sigma_{\text{exp}}$). No headers allowed. |
| `save-folder` | The program automatically creates an output directory named `RES_<save-folder>` in the current working path. |
| `qrange` | $q$-range to use for curve fitting: $[q_{\text{min}}, q_{\text{max}}]$. |
| `model` | Model class name (e.g., `LUV_POPC` or `LUV_POPE`). |
| `temperature-init` | Initial annealing temperature ($T_0$). Recommended $\approx 100 \times \chi_{\text{target}}^2$. |
| `target-X2` | $\chi_{\text{target}}^2$ value for adaptive convergence checking. |
| `data-binning` | Frequency of data points to keep for fast preliminary analyses. |
| `error-scale` | Scale factor $\in [0, 1]$ for $\sigma_{\text{exp}}$ for preliminary analyses (2% of $I_{\text{exp}}$ as lower limit for $\sigma_{\text{exp}}$). |
| `iterations` | Number of independent TSA runs ($N_I$). |
| `processes` | CPU cores used for parallel iteration. Set to 1 for serial mode. |
| `plot-only` | *on*: display data and initial guess curve only; *off*: run minimization algorithm. |

**Table 2.** List of positional arguments in the LUV_POPC model used as adjustable or fixed parameters (*y*) along with the associate Gaussian or non-informative priors (i.e., hand boundaries [*min, max*]).

| Model parameters |  | Prior initialization |  |  | Ref./Exp. |
| --- | --- | --- | --- | --- | --- |
| $y$ | Description | $\mu_y$ | $\varsigma_y/\mu_y$ | [$min, max$] | |
| $\mathcal{N}$ | scaling factor | $10^{-7\dagger}$ | - | - | - |
| $n_v$ | LUV concentration (number density $\text{\AA}^{-3}$) | - | - | [0.1, 2.5] | - |
| $R_v$ | LUV radius ($\text{\AA}$) | 450 | 0.10 | - | DLS exp.<sup>(a)</sup> |
| $Z$ | LUV polydispersity-related term | - | - | [5.1, 18] | - |
| $d_{\text{cholCH3}}$ | Relative position of group cholCH3 ($\text{\AA}$) | 4.3 | 0.03 | - | (Frewein et al., 2021) |
| $\delta_{\text{cholCH3}}$ | Fluctuation width of group cholCH3 ($\text{\AA}$) | 3.0 | 0.02 | - | (Kučerka et al., 2011) |
| $d_{\text{PCN}}$ | Relative position of group PCN ($\text{\AA}$) | 4.3 | 0.08 | - | (Frewein et al., 2021) |
| $\delta_{\text{PCN}}$ | Fluctuation width of group PCN ($\text{\AA}$) | 2.5 | 0.20 | - | (Frewein et al., 2021) |
| $d_{\text{CG}}$ | Relative position of group CG ($\text{\AA}$) | 0.8 | 0.08 | - | (Frewein et al., 2021) |
| $\delta_{\text{CG}}$ | Fluctuation width of group CG ($\text{\AA}$) | 2.5 | 0.20 | - | (Frewein et al., 2021) |
| $2D_C$ | Hydrophobic membrane thickness ($\text{\AA}$) | 28.4 | 0.03 | - | (Frewein et al., 2021) |
| $\sigma_C/2D_C$ | relative membrane thickness fluctuation | 0.079 | 0.06 | - | (Frewein et al., 2021) |
| $\delta_{\text{CH2}}$ | Fluctuation width of group CH<sub>2</sub> ($\text{\AA}$) | 2.48 | 0.05 | - | (Kučerka et al., 2011) |
| $d_{\text{CH}}$ | Position of group of group CH ($\text{\AA}$) | 9.0<sup>†</sup> | - | - | (Kučerka et al., 2011) |
| $\delta_{\text{CH}}$ | Fluctuation width of group CH ($\text{\AA}$) | 3.05<sup>†</sup> | - | - | (Kučerka et al., 2011) |
| $\delta_{\text{CH3}}$ | Fluctuation width of group CH<sub>3</sub> ($\text{\AA}$) | 3.40 | 0.05 | - | (Frewein et al., 2021) |
| $r_{\text{PCN}}$ | PCN relative head-group volume | 0.30 | 0.05 | - | (Kučerka et al., 2011) |
| $r_{\text{CG}}$ | CG relative head-group volume | 0.41 | 0.05 | - | (Kučerka et al., 2011) |
| $r_{12}$ | CH to CH<sub>2</sub> volume ratio | 0.80<sup>†</sup> | - | - | (Kučerka et al., 2011) |
| $r_{32}$ | CH<sub>3</sub> to CH<sub>2</sub> volume ratio | 1.93 | 0.05 | - | (Kučerka et al., 2011) |
| $T$ | Sample temperature ($^{\circ}\text{C}$) | 50<sup>†</sup> | - | - | - |
| $V_H$ | Lipid head-group volume ($\text{\AA}^3$) | 324 | 0.02 | - | (Nagle et al., 2019)<sup>(b)</sup> |
| $V_{\text{BW}}$ | Hydration-layer water volume ($\text{\AA}^3$) | 29.9 | 0.06 | - | (Frewein et al., 2021) |
| *Const.* | Constant background | - | - | $[3 \times 10^{-4}, 7 \times 10^{-4}]$ | - |

<sup>(a)</sup> Dynamic light scattering measurements (data not shown);
<sup>(b)</sup> Derived from the range of values reported for PC lipids;
<sup>†</sup> Fixed parameter.

Figure 3 shows the best fits for POPC, POPE, and DMPC-based LUV samples. The data are compared with model curves generated using the median values of each adjustable parameter distribution (Fig. 3A). These ‘median curves’ are in turn compared with the outcomes of individual iterations (Fig. 3B), highlighting which *q*-ranges are most affected by data uncertainties and parameter distribution widths. Additionally, the stochastic nature of the minimization algorithm is reflected in the 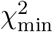 distribution profiles in Figure 3C. The adaptive convergence criteria enable a compact 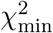 distribution, with a relative discrepancy between minimum and maximum 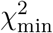 values of approximately 30% (POPC), 50% (POPE), and 20% (DMPC). In the cases of POPC and POPE, the large variations in *χ*^2^ values are primarily due to uncertainties in the LUV size (*R*_v_), as highlighted in the very low-*q* range in Figure 3B. In the case of POPE, the 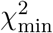 variability is further exacerbated by current model inconsistencies in describing the intensity minimum in the range *q* = 0.02–0.03 Å^−1^ (see, e.g., (Chappa et al., 2021)).

**Figure 3.**
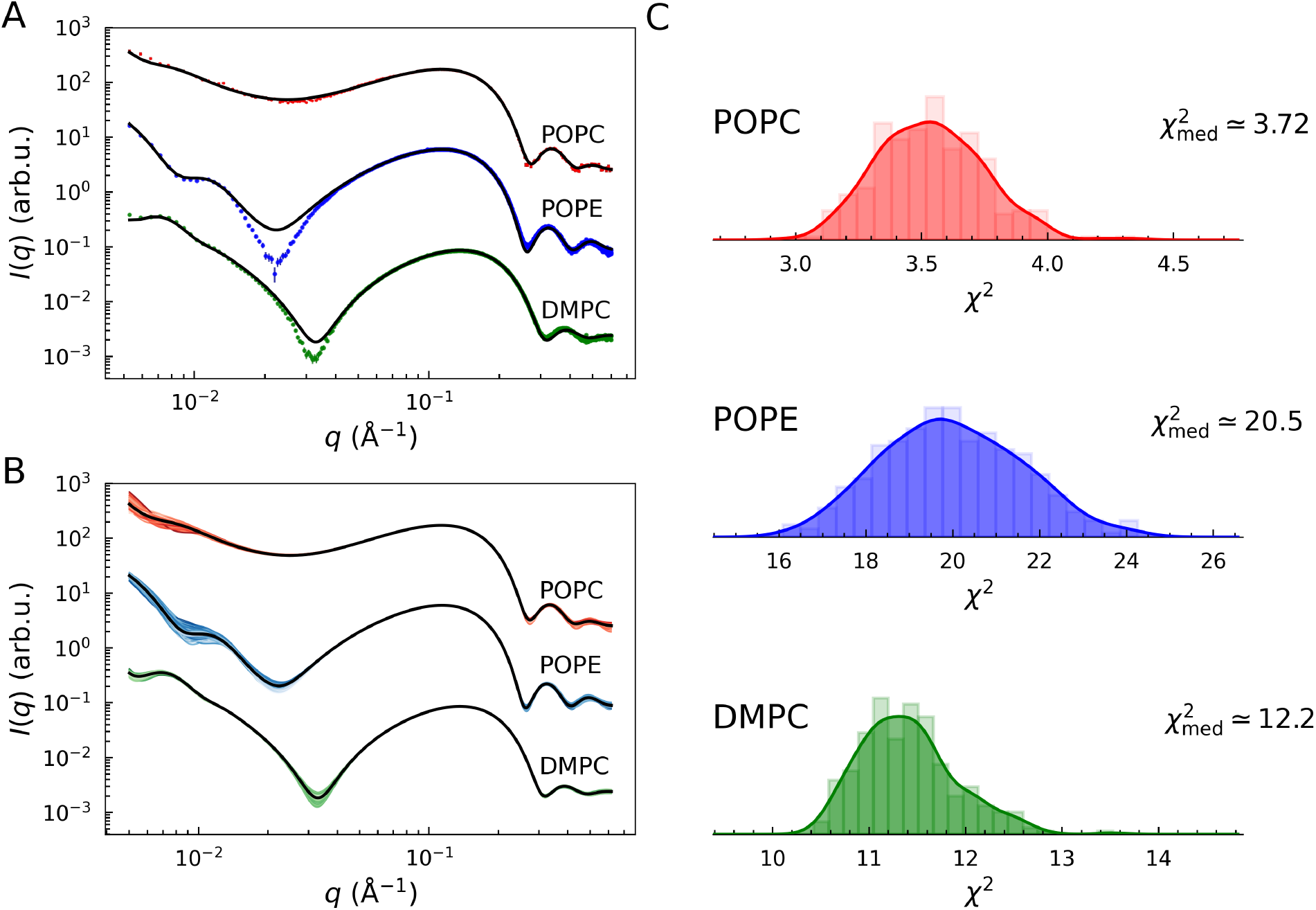
Best fits of POPC (red), POPE (blue), and DMPC (green) LUV SAXS data with *N*_I_ = 300. (A) Black lines display the model as a function of the parameter medians. These are compared to the best fits of each iteration in (B), where the shade of color reflects the best 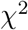 values (brighter for low values, darker for high values). (C) Distributions of 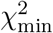 values. 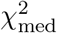 values were calculated using the adjustable parameter medians. Smooth curves display the kernel density estimates (KDEs) of the underlying histograms.

The results for the adjustable parameters are summarized in Table 3 for POPC LUVs, along with a comparison to previous results reported in (Kučerka et al., 2011) and (Frewein et al., 2021). The results and comparison for POPE and DMPC are provided in Tables S3 and S5. Note that our results are reported as median values and MADs. For reference, for a normal distribution, the relationship between standard deviation and MAD is *σ* ≃ MAD · 1.48.

**Table 3.** Results of the analysis of POPC at 50 ^°^C with the LUV_POPC model. Best values and relative uncertainties are reported as median `±` MAD (Median Absolute Deviation) and compared with previous results. Number of iterations *N*_*I*_ = 300. Note that relative uncertainties in previous works refer to standard deviations.

| Parameter $y$ | median±MAD | Ref. (Kučerka et al., 2011) | Ref. (Frewein et al., 2021) |
| --- | --- | --- | --- |
| $R_v$ (Å) | $440 \pm 30$ (7%) | n.a. | n.a. |
| $Z$ | $9.8 \pm 1.3$ (13%) | n.a. | n.a. |
| $d_{\text{CholCH}_3}$ (Å) | $4.29 \pm 0.10$ (2%) | $1.0^\ddagger$ | $4.3^\ddagger$ |
| $\delta_{\text{CholCH}_3}$ (Å) | $3.00 \pm 0.04$ (2%) | $2.98^\dagger$ | $3.00^\dagger$ |
| $d_{\text{PCN}}$ (Å) | $4.2 \pm 0.3$ (6%) | $4.5^\ddagger$ | $4.1^\ddagger$ |
| $\delta_{\text{PCN}}$ (Å) | $2.6 \pm 0.3$ (10%) | $2.81$ (2%) | $2.5$ (20%) |
| $d_{\text{CG}}$ (Å) | $0.81 \pm 0.04$ (5%) | $0.75^\ddagger$ | $0.80^\ddagger$ |
| $\delta_{\text{CG}}$ (Å) | $2.2 \pm 0.2$ (9%) | $2.48$ (2%) | $2.5$ (20%) |
| $2D_C$ (Å) | $28.2 \pm 0.4$ (2%) | $28.1^\ddagger$ (2%) | $28.4^\ddagger$ (3%) |
| $\varepsilon_C = \sigma_C/2D_C$ | $0.076 \pm 0.002$ (3%) | n.a. | $0.079$ (6%) |
| $\delta_{\text{CH}_2}$ (Å) | $2.49 \pm 0.09$ (4%) | $2.50^{(s)}$ (2%) | $2.5^\dagger$ |
| $d_{\text{CH}}$ (Å) | $9^\dagger$ | $9^\dagger$ | $9^\dagger$ |
| $\delta_{\text{CH}}$ (Å) | $3.05^\dagger$ | $3.05^\dagger$ | $3.05^\dagger$ |
| $\delta_{\text{CH}_3}$ (Å) | $3.40 \pm 0.13$ (4%) | $2.69$ (2%) | $3.40$ (5%) |
| $r_{\text{PCN}}$ | $0.312 \pm 0.012$ (4%) | $0.300^{(s)}$ (2%) | $0.29^\dagger$ |
| $r_{\text{CG}}$ | $0.433 \pm 0.015$ (4%) | $0.41^{(s)}$ (2%) | $0.45^\dagger$ |
| $r_{12}$ | $0.80^\dagger$ | $0.80^{(s)}$ (2%) | $0.80^\dagger$ |
| $r_{32}$ | $1.93 \pm 0.07$ (4%) | $1.93^{(s)}$ (2%) | $2.09^\dagger$ |
| $T$ (°C) | $50^\dagger$ | - | - |
| $V_H$ (Å<sup>3</sup>) | $323 \pm 5$ (2%) | $331^\dagger$ | $320^\dagger$ |
| $V_{\text{BW}}$ (Å<sup>3</sup>) | $30.43 \pm 0.04$ (0.13%) | n.a. | $29.9$ (6%) |
| **Calculated parameters** |  |  |  |
| $z_{\text{CholCH}_3}$ (Å) | $23.4 \pm 1.4^\ddagger$ (6%) | $20.3$ (2%) | $23.4$ (3%) |
| $z_{\text{PCN}}$ (Å) | $19.1 \pm 0.4^\ddagger$ (2%) | $19.3$ (2%) | $19.1$ (8%) |
| $z_{\text{CG}}$ (Å) | $14.9 \pm 0.3^\ddagger$ (2%) | $14.8$ (2%) | $15.0$ (8%) |
| $V_L$ (Å<sup>3</sup>) | $1274.85^\ddagger$ | $1275.5^\dagger$ | $1276.9^\dagger$ |
| $a_L$ (Å<sup>2</sup>) | $67.4 \pm 1.0^\ddagger$ (1.5%) | $67.3$, (2%) | $67.5$ (2%) |
| $D_B$ (Å) | $37.8 \pm 0.6^\ddagger$ (1.5%) | $37.9^\ddagger$ (2%) | $38.4^\ddagger$ (5%) |
| $D_{\text{pp}}$ (Å) | $36.7^\ddagger$ (2%) | $35.9^\ddagger$ (2%) | $37.5^\ddagger$ (3%) |
| $D_{\text{H}_1}$ (Å) | $4.8 \pm 0.2^\ddagger$ (4%) | $3.91^\ddagger$ (2%) | $4.6^\ddagger$ (20%) |
| $n_w$ | $\sim 17^\ddagger$ | n.a. | $14^\ddagger$ (6%) |

$^\dagger$ fixed values; $^\ddagger$ calculated values; $^{(s)}$ ‘soft’ Bayesian constraints (Kučerka et al., 2011);

It is important to note a methodological difference when comparing our results with the reference studies by (Kučerka et al., 2011) (PC species) and (Kučerka et al., 2015) (PE species). In those works, vesicles were prepared by extrusion through 50 nm filters, yielding small unilamellar vesicles (SUVs), whereas our analysis is based on 100 nm LUVs. The higher membrane curvature of SUVs compared to LUVs can induce a weak but systematic membrane thinning (Chappa et al., 2021). In addition, the reference systems were analyzed over a more restricted *q*-range (*q*_min_ ~ 0.05 Å^−1^) using a joint SAXS/SANS analysis approach. Despite these differences in vesicle size, *q*-range, and overall analysis methodology our results demonstrate remarkable agreement with the earlier findings. This consistency suggests, on the one hand, that curvature effects are negligible for POPC and POPE under the conditions considered, and, on the other hand, highlights the robustness of the *SAS_MoCa* minimization engine in delivering reliable parameter estimates and confidence intervals, even with a relatively large number of adjustable parameters.

For DMPC, we faced a specific challenge regarding prior knowledge. While structural parameters for POPC and POPE lipids are well-documented in the literature, information regarding the hydration layer volume, *V*_BW_, and the acyl-chain thickness fluctuation width, *σ*_C_, for DMPC was unavailable.

Consequently, for the DMPC analysis, we adopted broad, non-informative rectangular priors (hard boundaries) for *V*_BW_ and *σ*_C_. Remarkably, even without informative priors for these parameters, the TSA algorithm converged to physically meaningful values (Tab. S5), demonstrating the robustness of the minimization engine even for under-constrained parameters.

In addition to reporting point estimates, the full strength of the Bayesian framework implemented in *SAS_MoCa* lies in inspecting the posterior distributions of the adjustable parameters, their correlations, and their relationship to the chosen priors. Figure 4 shows the posterior probability densities of selected parameters together with their corresponding prior distributions. These posterior distributions constitute the primary output of the program and should always be examined before interpreting the associated median and MAD values. Parameters in Figs. 4A and B illustrate cases where the posteriors remain closely aligned with their priors. By contrast, the distributions in Figs. 4C, D, and F exhibit a clear shift in center of mass relative to the prior mean, exemplifying data-driven updating of informative priors. Figure 4E shows a similar data-driven update, but starting from a non-informative prior. The full set of posterior results for all adjustable parameters is provided in Figs. S2–S4.

**Figure 4.**
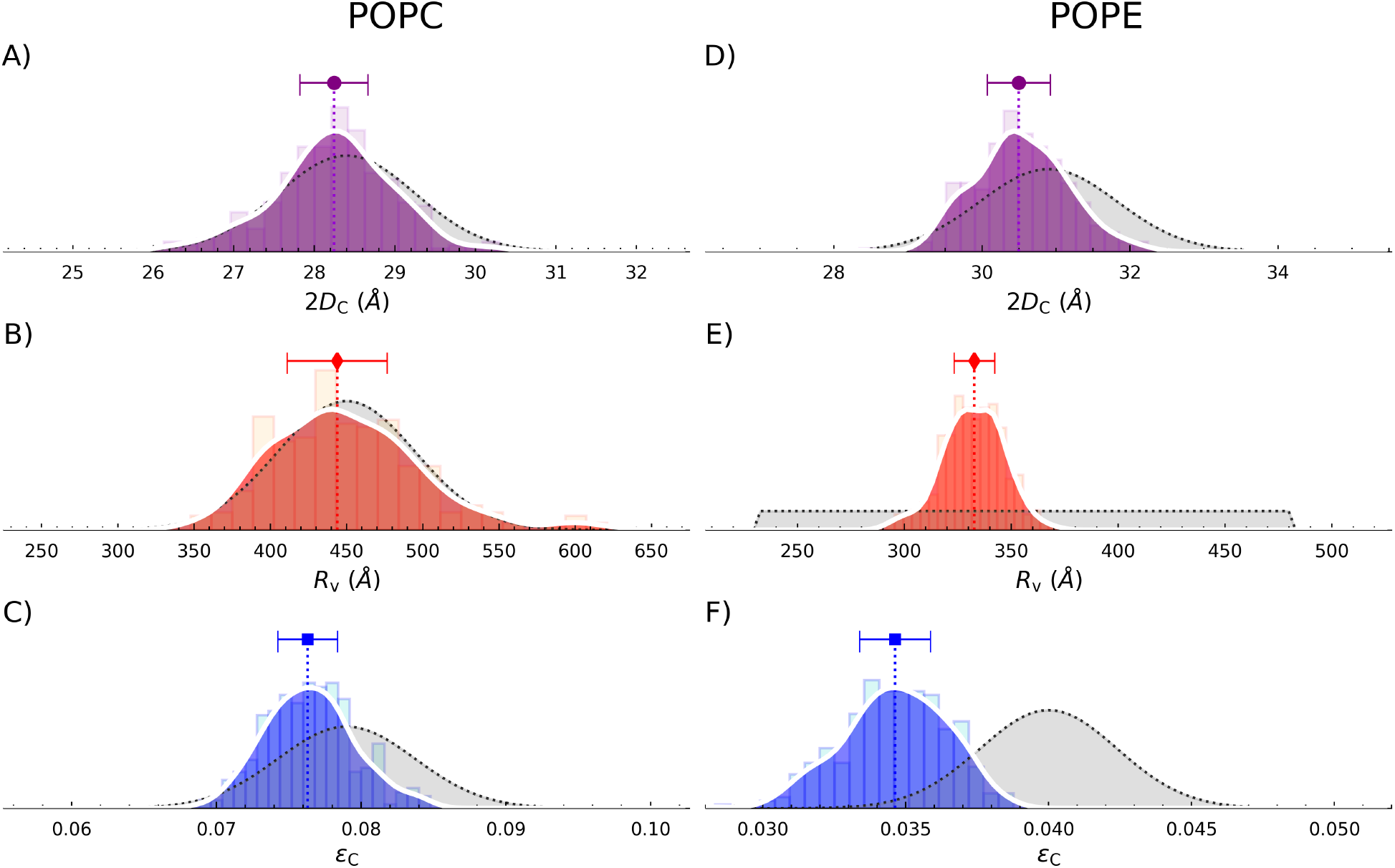
Comparison between probability density distributions (posteriors) and priors of three selected adjustable parameters obtained during the curve fitting of POPC (A-C) and POPE systems (D-F). (A,D) Acyl-chain thickness 2*D*_C_; (B,E) vesicle radius *R*_v_; and (C,F) relative variation of acyl-chain thickness *ε*_C_ = *σ*_C_*/*2*D*_C_. Histograms contain *N*_I_ = 300 entries; colored curves represent KDEs, while dotted gray lines show the associated prior distribution. Dots with error bars highlight the median and MAD values for each probability distribution.

Based on the found posterior distributions the program automatically generates a heatmap representing the matrix of Pearson correlation coefficients for the adjustable parameters. Figure 5 shows a selected example of the correlation between adjustable parameters. Full correlation matrices for POPC, POPE, and DMPC systems are shown in Figs. S5–S7.

**Figure 5.**
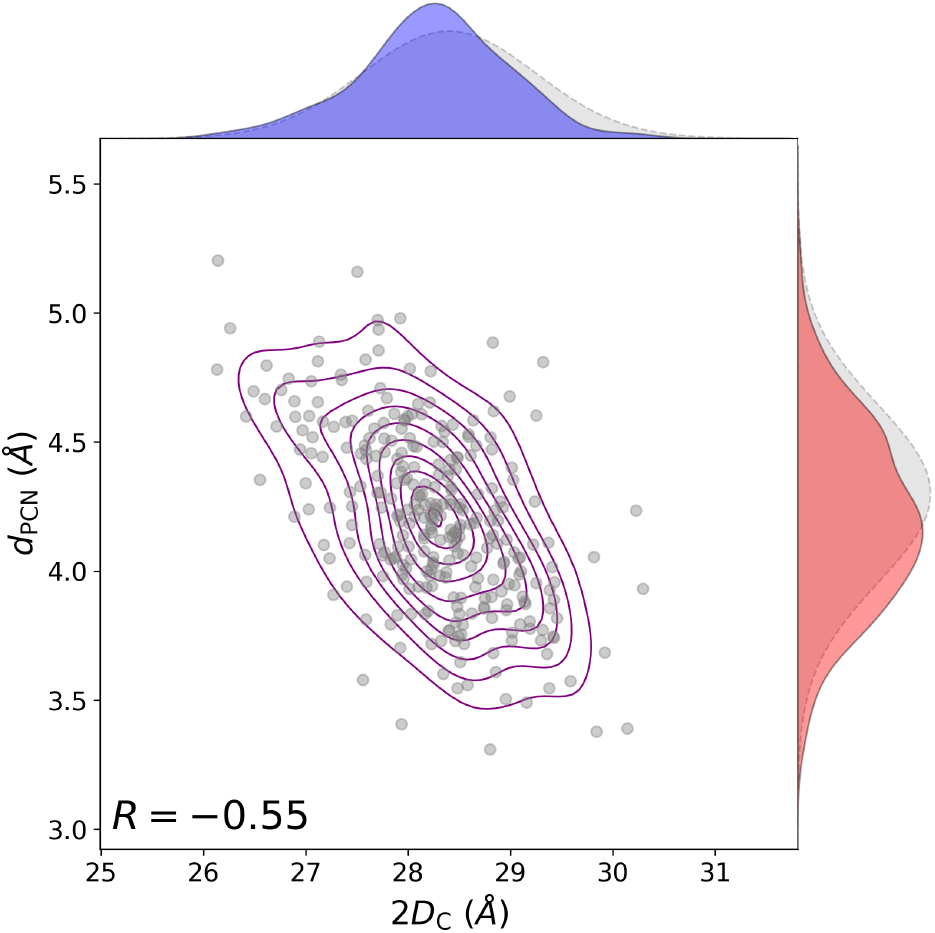
KDE bivariate density map and associated probability densities of two selected adjustable parameters obtained during the curve fitting of POPC systems: acyl-chain thickness 2*D*_C_ (blue, *x*-axis) and distance between PCN and CG quasi-molecular groups *R*_PCN_ (red, *y*-axis). Results from *N*_I_ = 300 iterations. Dashed gray lines indicate the associated prior distribution. *R* is the Pearson correlation coefficient.

### 4.2 Benchmarking Parameter Sensitivity and Prior Choices

We investigated how priors affect both sensitive and non-sensitive adjustable parameters. To this end, we reanalyzed the POPC data while systematically modifying the prior *π*(*µ*_*y*_, 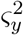) for one parameter at a time. Specifically, we shifted the prior mean *µ*_*y*_ in steps of ±*ς*_*y*_ and examined the resulting posterior distribution of the corresponding parameter *y*.

As illustrated in Fig. 6 for two representative parameters, we observed two characteristic behaviors. Figure 6A shows a sensitive parameter, the acyl chain thickness 2*D*_C_. Here, the posterior is dominated by the likelihood and remains largely unchanged when the prior is shifted, indicating that the data strongly constrain 2*D*_C_. In contrast, Fig. 6B shows a non-sensitive parameter, the lipid headgroup volume *V*_H_. In this case, the posterior distributions closely track the shifted priors, demonstrating that the posterior is prior-dominated and essentially determined by the initial belief—even when that belief corresponds to non-physical values.

**Figure 6.**
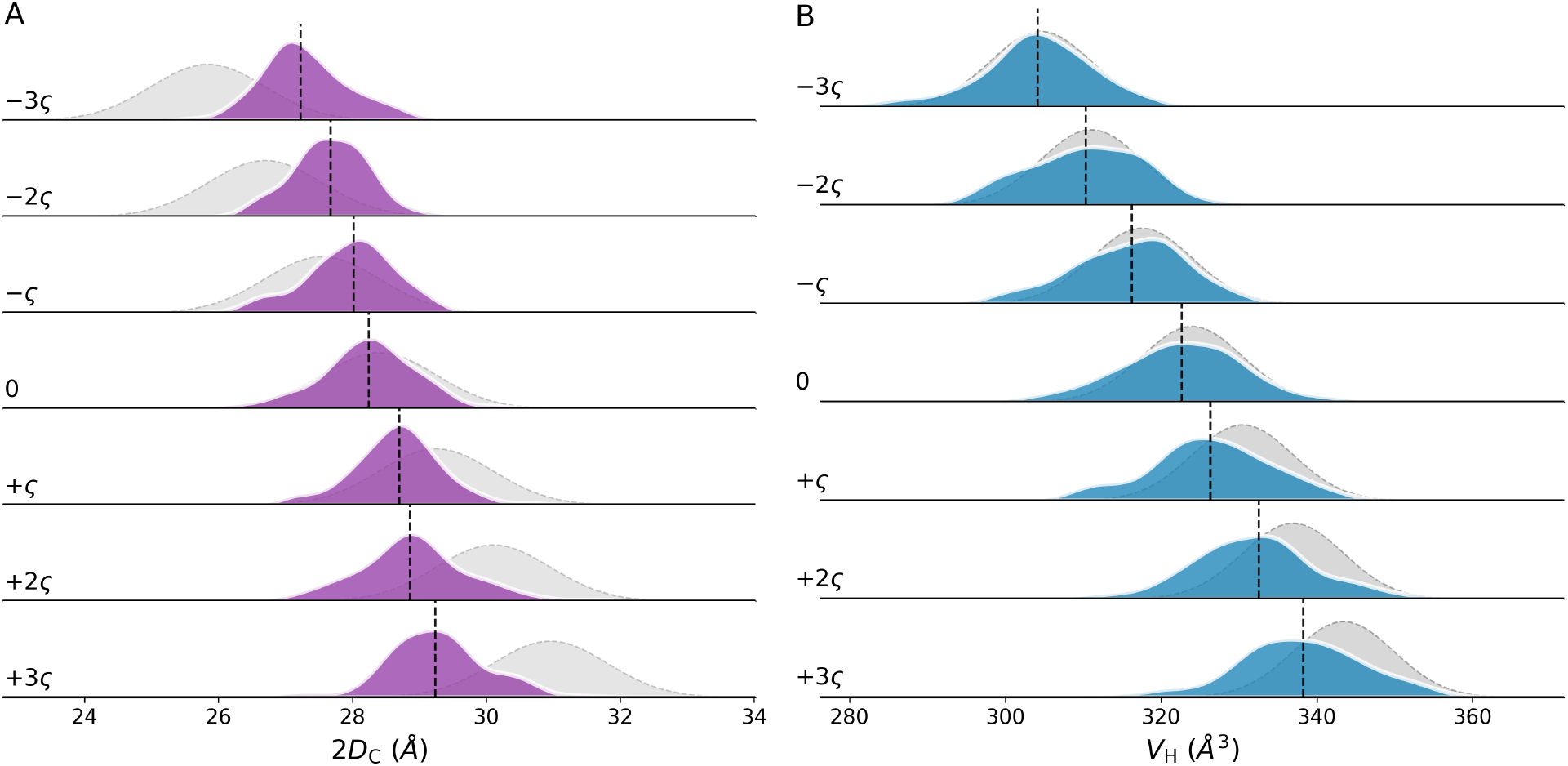
Verifying the dependency of parameter distributions on priors: sensitive vs. non-sensitive parameters. (A) Acyl chain thickness 2*D*_C_ distributions (KDE representation in purple) against priors shifted by ±*nς*_y_ (gray). (B) Lipid head-group volume *V*_H_ distributions (KDE representation in blue) against priors shifted by ±*nς*_y_ (gray). Vertical dashed lines indicate the medians of the adjustable parameter distributions. Number of iterations for each independent analysis *N*_I_ = 100.

It is worth emphasizing that even for sensitive parameters such as 2*D*_C_, the choice of prior can still induce a noticeable shift in the posterior median. Users should therefore exercise care when specifying priors: the principle of “garbage in, garbage out” applies, and the reliability of the inferred parameters depends critically on the quality and physical plausibility of the prior information.

### 4.3 Reproducibility and Practical Guidelines

Accessing the sampling distributions of each adjustable parameter enables the application of basic statistical inference tools. In particular, we can utilize these distributions to verify the consistency of results across independent analyses performed with varying numbers of iterations, *N*_I_. For normal-like (i.e., symmetric, bell-shaped) distributions, the median is comparable to the mean, 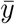, and the standard deviation is approximated by *σ*_*y*_ ≃ 1.48 · MAD. Consequently, one can estimate the confidence intervals (CIs) for both 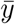 and *σ*_*y*_ using the pivotal method (Ramachandran & Tsokos, 2015). Specifically, the (1 − *α*) · 100% CIs for the mean 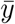 as *N*_I_ → ∞ are given by:

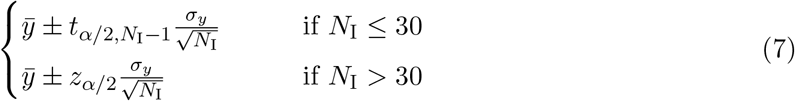

where *α* is the significance level, 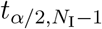 represents the critical values of the Student’s *t*-distribution with *N*_I_ −1 degrees of freedom, and *z*_*α/*2_ denotes the corresponding quantiles of the standard normal distribution for large sample sizes (Ramachandran & Tsokos, 2015).

Similarly, the (1 − *α*) · 100% CIs for the variance 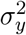 are:

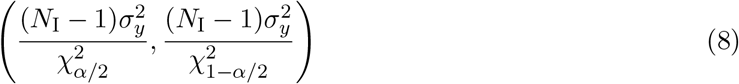

where 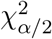 are the critical values of the *χ*^2^-distribution tails (Ramachandran & Tsokos, 2015).

Figure 7 illustrates the reproducibility of results for varying *N*_I_ and compares the empirical distributions with predicted CIs. The latter are calculated by averaging the 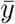 and *σ*_*y*_ outcomes for *N*_I_ = 300 as references for Eqs. 7 and 8. As shown in Fig. 7A, the distributions of the medians are well-contained within the predicted confidence intervals. However, for the standard deviations (Fig. 7B), the empirical spread appears larger than the theoretical prediction, particularly for small *N*_I_. This is likely due to the fact that, e.g., the posterior distributions are not perfectly normal.

**Figure 7.**
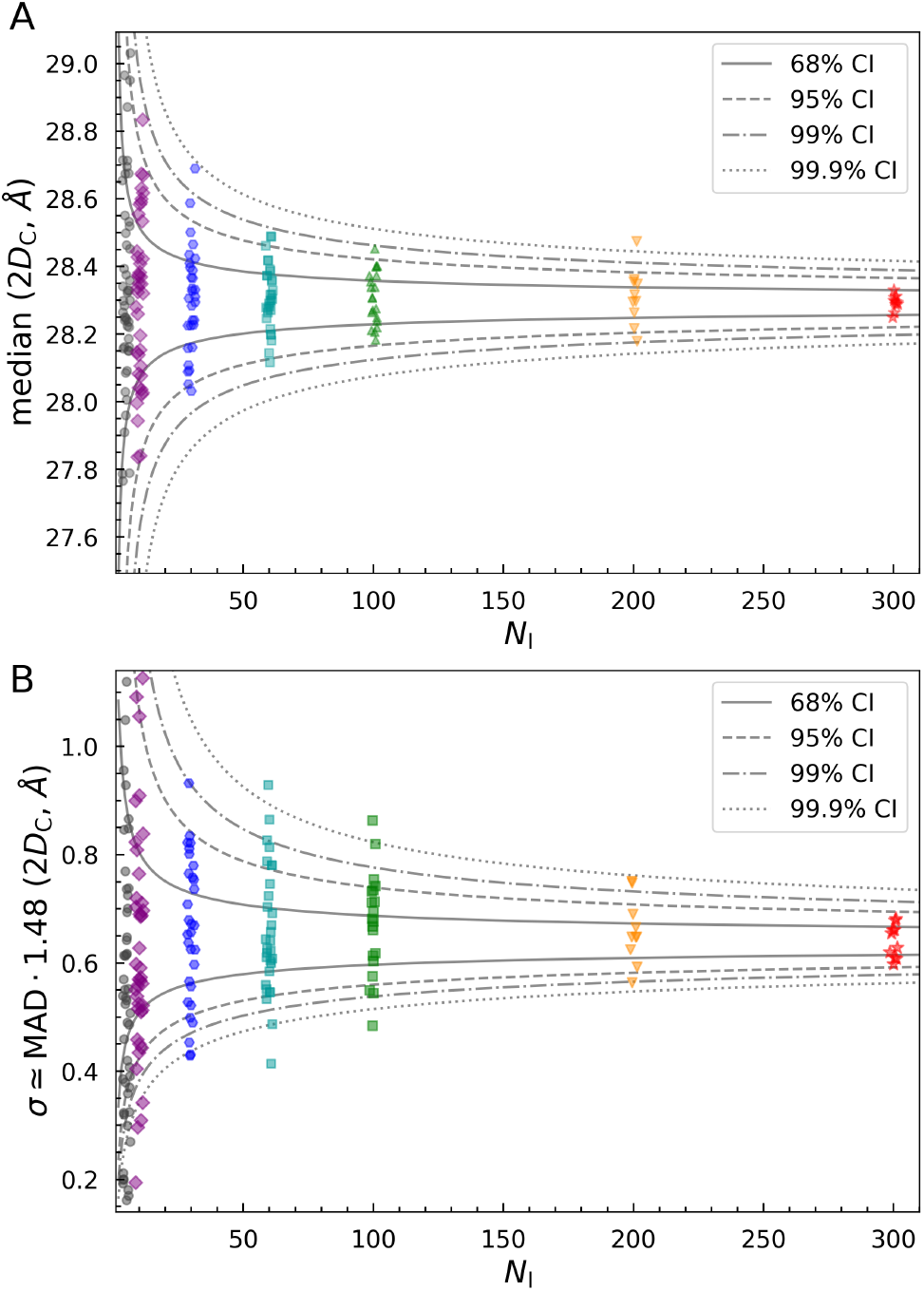
Median (A) and equivalent standard deviation (B) 2*D*_C_ values from independent analyses performed at different *N*_I_ values. Lines mark the ranges for different levels of CIs. These are calculated using Eqs. 7 (A) and 8 (B), by using the average of the outcomes for *N*_I_ = 300 as 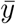 and *σ*_*y*_ values.

These reproducible behaviors in
form a recommended workflow for *SAS_MoCa* users:

1. **Exploratory Phase:** Perform rapid tests with a small number of iterations (*N*_I_ ≈ 5 − 20) and varied initializations to assess parameter sensitivity, validate posterior results and refine prior choices.
2. **Precision Planning:** Invert Eqs. 7 and 8 and use preliminary results to estimate the required *N*_I_ to achieve a target confidence interval width, optimizing computational resources.
3. **Conclusive Analysis:** Execute a computationally intensive run with, e.g., *N*_I_ *>* 200 to generate robust posterior distributions, once the model initialization is validated.

This tiered approach ensures that computational effort is allocated efficiently, balancing rapid feedback with statistical rigor.

## 5 Conclusion

We have presented *SAS_MoCa*, an open-source computational framework designed to address a critical bottleneck in the structural characterization of LUVs via small-angle X-ray scattering (SAXS). To our knowledge, *SAS_MoCa* represents the first freely available software capable of performing compositional scattering density profile (SDP) analysis tailored specifically for LUVs, filling a significant gap left by discontinued proprietary tools and general-purpose packages lacking specialized workflows for membrane compositional modeling.

The core innovation of *SAS_MoCa* lies in its rigorous application of Bayesian inference to SAXS data analysis. By integrating adaptive thermodynamic simulated annealing with flexible prior probability distributions, the software enables robust statistical quantification of parameter uncertainties and correlations, even in highly under-constrained regimes typical of single-curve scattering experiments. This approach fundamentally shifts the analytical workflow from heuristic trial-and-error fitting to transparent statistical inference. Users are no longer forced to accept point estimates without context; instead, the explicit comparison of prior and posterior distributions provides a diagnostic view of which structural parameters are truly resolved by the experimental data and which remain constrained by external knowledge.

Our validation across three distinct lipid systems—POPC, POPE, and DMPC—demonstrated that *SAS_MoCa* yields results consistent with established joint SAXS/SANS literature, despite analyzing single-curve datasets with significantly higher degrees of freedom. Crucially, the benchmarking of parameter sensitivity highlights the necessity of physically meaningful priors: while the minimization engine is robust, the reliability of the posterior remains bounded by the validity of the initial assumptions. This feature enforces a higher standard of scrutiny, requiring researchers to explicitly justify their structural hypotheses rather than relying on implicit constraints within black-box algorithms.

Looking ahead, the modular architecture of *SAS_MoCa* is designed for extensibility. Current development efforts aim to incorporate multiscale models for proteoliposomes and extend the formalism to handle small-angle neutron scattering (SANS), including joint contrast-variation analyses. Furthermore, we plan to expand the library of compositional models to cover asymmetric LUVs (aLUVs) and bacterial systems (Semeraro et al., 2017; Semeraro et al., 2021) and minimal cells, thereby broadening the applicability of the framework to increasingly complex biomimetic membrane systems.

*SAS_MoCa* is released under the BSD 3-Clause license and is publicly available at https://github.com/PabstLab/SAS_MoCa (Semeraro & Pabst, 2026). We envision *SAS_MoCa* democratizing high-resolution membrane structural analysis—empowering laboratories to perform rigorous, compositional investigations with full statistical transparency. By making advanced Bayesian inference accessible to the broader biophysics community, the tool fosters a culture of reproducible reporting where assumptions are explicit and uncertainties are quantified.

## A Scattering intensity

### A.1 Vesicle size and polydispersity: separated form factor model

In the SFF framework the LUVs shape is described by an infinitesimally thin spherical frame of radius *R*_*v*_. The radius polydispersity is described by the Schulz probability distribution function, that ensures an analytical form (Pencer et al., 2006). The scattering intensity of such a twodimensional, polydisperse set of spherical frames is given by *P* (*q, R*_*v*_) = 16*π*^2^*µ*_4_(*qR*_*v*_), where

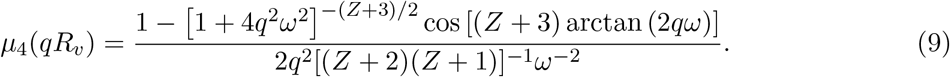

Here 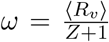, and *Z* is the width parameter of the Schulz distribution (the variance is defined as *σ*^2^ = ⟨*R*_*v*_⟩^2^*/*(*Z* + 1)). Note that 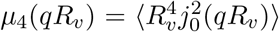, where *j*_0_(*qR*_*v*_) is the spherical Bessel function of 0’th order.

The limit for zero scattering angle (forward scattering) is

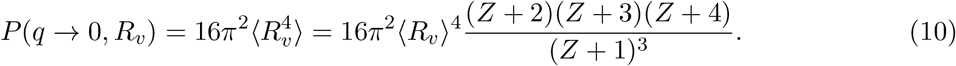

### A.2 Transbilayer structure: scattering density profile

The scattering amplitude for the transbilayer structure of lipid bilayers, *A*_M_(*q*), refers to the latest update of the SDP model (Frewein et al., 2021). The SDP approach is a compositional model that takes into account the chemical compositions of quasi-molecular groups in which the lipids are parsed. Therefore, the specific scattering amplitude contributions depends on the lipid composition. In the following, we describe the basic models pertaining to the quasi-molecular parsing of POPC, POPE and DMPC, as described in to (Kučerka et al., 2011; Kučerka et al., 2015; Frewein et al., 2021).

The base scattering amplitude is given by:

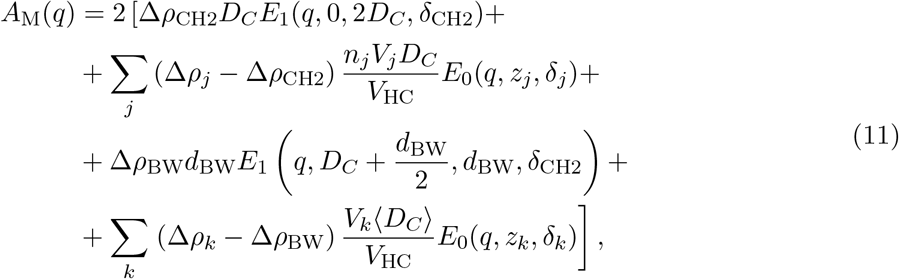

where the functions

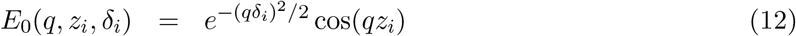

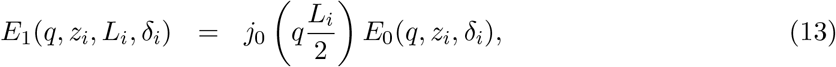

are the Fourier transform of a Gaussian profile (12) and a error-function-based smooth rectangular profile (13). The subscript *j* refers to the chain quasi-molecular group CH3 and CH; the subscript *k* includes the lipid head groups (e.g., CG, PCN, and cholCH3 in the case of POPC, see Tab. S1). For the generic *i*-th quasi-molecular group the parameter *z*_*i*_ is the center of the Gaussian or smooth rectangular functions; *δ*_*i*_ is the associated width of either the Gaussian profile or the error functions composing the smooth rectangular profile. In the latter case the distance between the centers of the two error functions is *L*_*i*_.

The acyl-chain thickness fluctuation average, ⟨· · · ⟩_HC_, is accounted for with the expression

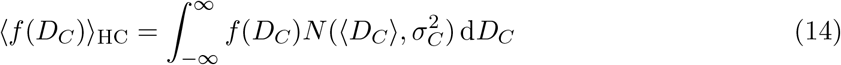

where *N* (⟨*D*_*C*_⟩, 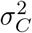) is the Gaussian probability distribution function of mean ⟨*D*_*C*_⟩ and standard deviation *σ*_*C*_ (Frewein et al., 2021).

The parameters describing *A*_M_ are described and listed in Table 4. For details refer to (Frewein et al., 2021) and (Kučerka et al., 2008).

**Table 4.** Recap of the structural parameters of interest relative to *A*_M_(*q*) (Kučerka et al., 2008; Frewein et al., 2021).

| Description | Parameter |
| --- | --- |
| SLD of group $x$ | $\rho_i$ |
| SLD of medium | $\rho_{\text{sol}}$ |
| SLD contrast of group $x$ | $\Delta\rho_i = \rho_i - \rho_{\text{sol}}$ |
| Relative position of group $i$ | $d_i$ |
| Fluctuation width of group $i$ | $\delta_i$ |
| Molecular volume of group $i$ | $V_i$ |
| Number of group $i$ units | $n_i$ |
| Width of hydration layer | $d_{\text{BW}} = 3.1 \text{ \AA}$ |
| Lipid volume | $V_L$ |
| Lipid head-group volume | $V_H$ |
| Lipid acyl-chain volume | $V_{\text{HC}} = V_L - V_H$ |
| Hydrophobic thickness | $2D_C$ |
| Acyl-chain thickness fluctuation width | $\sigma_C$ |
| Lateral area per lipid | $A_L = V_{\text{HC}}/D_C$ |
| Luzzati bilayer thickness | $D_B = 2V_L/A_L$ |
| Phosphate-phosphate distance | $D_{\text{pp}}$ |
| CH<sub>3</sub> to CH<sub>2</sub> volume ratio | $r_{32} = V_{\text{CH3}}/V_{\text{CH2}}$ |
| CH to CH<sub>2</sub> volume ratio | $r_{12} = V_{\text{CH}}/V_{\text{CH2}}$ |
| $k$-th group rel. volume | $r_k = V_k/V_H$ |
| Hydration-layer water volume | $V_{\text{BW}}$ |

## Acknowledgements

We would like to thank the BM29 beamline staff at the ESRF (Grenoble, France) for support during beamtime (proposal No. MX-2282). We thank Paulina Piller for the help in sample preparation; Moritz Frewein for the fruitful discussion; and Shahrzad Dabironezare for testing the repository and identifying bugs.

## Funding

This project has received funding from the European Union’s Horizon Europe research and innovation programme under grant agreement No 101169269. This research was funded in whole or in part by the Austrian Science Fund (FWF) [10.55776/PIN7380924].

### Data availability

Data and example scripts used in this study are openly available in the `examples` directory of the SAS_MoCa GitHub repository: https://github.com/PabstLab/SAS_MoCa.

