## SUPPLEMENTARY INFORMATION for "*SAS_MoCa*: a software for small-angle scattering data analysis of large unilamellar vesicles"

### S1 Lipids Quasi-Molecular Parsing

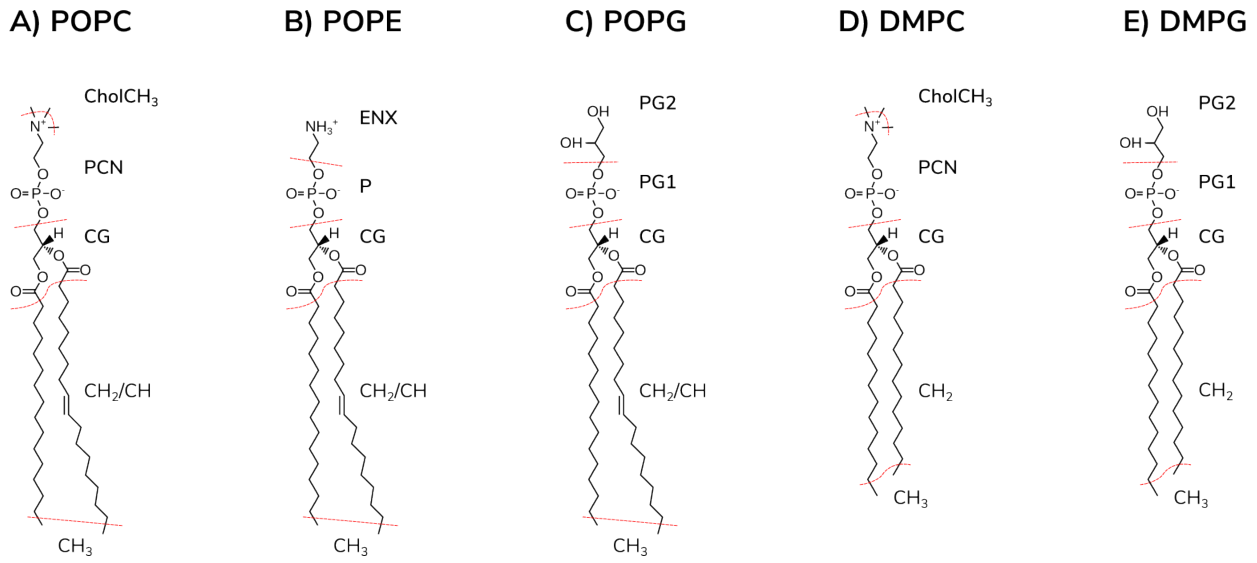


Figure S1: Chemical structures of the presently studied phospholipids and their parsing into quasi-molecular fragments.

Table S1: List of quasi-molecular groups for PC, PE and PG lipids, along with abbreviation, composition and X-ray scattering length $b$ (Kučerka et al., 2011; Kučerka et al., 2015; Pan et al., 2012).

| **Quasimolecular group** | **Abbreviation** | **Composition** | $b \times 10^{-4}$ (Å) |
|---|---|---|---|
| Terminal methyl group | CH3 | $CH_3$ | 2.53615 |
| Methylene group | CH2 | $CH_2$ | 2.25435 |
| Methine group | CH | CH | 1.97256 |
| Carbonyl–glycerol backbone | CG | $C_5H_5O_4$ | 18.8802 |
| Phosphate group + CN | PCN | $C_2H_4NO_4P$ | 19.7256 |
| Phosphate group | P/PG1 | $O_4P$ | 13.2443 |
| Choline-CH3 group | cholCH3 | $C_3H_9$ | 7.60844 |
| Amine group | ENX | $C_2H_7N$ | 7.32664 |
| Glycerol group | PG2 | $C_3H_7O_2$ | 11.5536 |
| Hydration layer | BW | $H_2O$ | 2.8179 |

### S2 Prior Strategies and Results

Table S2: List of positional arguments in the `LUV_POPE` model used as adjustable or fixed parameters ($y$) along with the associate Gaussian or non-informative priors (i.e., hand boundaries $[min, max]$).

| **Model parameters** | | **Prior initialization** | | | **Ref./Exp.** |
|---|---|---|---|---|---|
| $y$ | Description | $\mu_y$ | $\varsigma_y/\mu_y$ | $[min, max]$ | |
| $\mathcal{N}$ | scaling factor | $10^{-7}$† | - | - | - |
| $n_{\mathrm{v}}$ | LUV concentration (number density Å$^{-3}$) | - | - | $[0.1, 10]$ | - |
| $R_{\mathrm{v}}$ | LUV radius (Å) | - | - | $[230, 480]$ | - |
| $Z$ | LUV polydispersity-related term | - | - | $[8, 40]$ | - |
| $d_{\mathrm{ENX}}$ | Relative position of group ENX (Å) | 3.4 | 0.03 | - | (a) |
| $\delta_{\mathrm{ENX}}$ | Fluctuation width of group ENX (Å) | 2.8 | 0.2 | - | (a) |
| $d_{\mathrm{P}}$ | Relative position of group P (Å) | 3.1 | 0.08 | - | (a) |
| $\delta_{\mathrm{P}}$ | Fluctuation width of group P (Å) | 2.27 | 0.2 | - | (a) |
| $d_{\mathrm{CG}}$ | Relative position of group CG (Å) | 0.9 | 0.08 | - | (a) |
| $\delta_{\mathrm{CG}}$ | Fluctuation width of group CG (Å) | 2.27 | 0.20 | - | (a) |
| $2D_{\mathrm{C}}$ | Hydrophobic membrane thickness (Å) | 30.9 | 0.03 | - | (a) |
| $\sigma_{\mathrm{C}}/2D_{\mathrm{C}}$ | relative membrane thickness fluctuation | 0.04 | 0.06 | - | (Frewein et al., 2023) |
| $\delta_{\mathrm{CH2}}$ | Fluctuation width of group $CH_2$ (Å) | 2.62 | 0.05 | - | (a) |
| $d_{\mathrm{CH}}$ | Position of group of group CH (Å) | 9.43 | 0.02 | - | (Kučerka et al., 2015) |
| $\delta_{\mathrm{CH}}$ | Fluctuation width of group CH (Å) | 2.8† | - | - | (Kučerka et al., 2015) |
| $\delta_{\mathrm{CH3}}$ | Fluctuation width of group $CH_3$ (Å) | 2.81 | 0.05 | - | (a) |
| $r_{\mathrm{P}}$ | P relative head-group volume | 0.14 | 0.05 | - | (Kučerka et al., 2015) |
| $r_{\mathrm{CG}}$ | CG relative head-group volume | 0.55 | 0.05 | - | (Kučerka et al., 2015) |
| $r_{12}$ | CH to $CH_2$ volume ratio | 0.80 | 0.05 | - | (Kučerka et al., 2015) |
| $r_{32}$ | $CH_3$ to $CH_2$ volume ratio | 1.95 | 0.05 | - | (Kučerka et al., 2015) |
| $T$ | Sample temperature (°C) | 50† | - | - | - |
| $V_{\mathrm{H}}$ | Lipid head-group volume (Å$^3$) | 247 | 0.02 | - | (a) |
| $V_{\mathrm{BW}}$ | Hydration-layer water volume (Å$^3$) | 29.6 | 0.06 | - | (Frewein et al., 2023) |
| *Const.* | Constant background | - | - | $[\times 10^{-3}, 2.2 \times 10^{-3}]$ | - |

(a) Combined from (Kučerka et al., 2015) and (Frewein et al., 2023);
† Fixed parameter.

Table S3: Results of the analysis of POPE at 50 °C with the LUV_POPE model. Best values and relative uncertainties are reported as median ± MAD (Median Absolute Deviation) and compared with previous results. Number of iterations $N_I = 300$. Note that relative uncertainties in previous works refer to standard deviations.

| **Parameter** $y$ | median±MAD | Ref. (Kučerka et al., 2015) | Ref. (Frewein et al., 2023) |
|---|---|---|---|
| $R_{\rm v}$ (Å) | $330 \pm 10\,(3\%)$ | n.a. | n.a. |
| $Z$ | $21 \pm 3\,(17\%)$ | n.a. | n.a. |
| $d_{\rm ENX}$ (Å) | $3.40 \pm 0.09\,(3\%)$ | $1.74^{\ddagger}$ | $5.0^{\ddagger}$ |
| $\delta_{\rm ENX}$ (Å) | $3.1 \pm 0.5\,(16\%)$ | $2.8^{\dagger}$ | $2.8^{\dagger}$ |
| $d_{\rm P}$ (Å) | $3.16 \pm 0.19\,(6\%)$ | $4.33^{\ddagger}$ | $1.8^{\ddagger}$ |
| $\delta_{\rm P}$ (Å) | $2.77 \pm 0.17\,(6\%)$ | 2.27 (2%) | 3.5 (20%) |
| $d_{\rm CG}$ (Å) | $0.90 \pm 0.07\,(8\%)$ | $0.30^{\ddagger}$ | $1.4^{\ddagger}$ |
| $\delta_{\rm CG}$ (Å) | $2.2 \pm 0.2\,(9\%)$ | 2.27 (2%) | 2.5 (20%) |
| $2D_{\rm C}$ (Å) | $30.5 \pm 0.4\,(1.4\%)$ | $30.8^{\ddagger}$ (2%) | $31.2^{\ddagger}$ (3%) |
| $\varepsilon_{\rm C}$ | $0.0346 \pm 0.0012\,(4\%)$ | n.a. | 0.04 (6%) |
| $\delta_{\rm CH2}$ (Å) | $2.63 \pm 0.11\,(4\%)$ | $2.62^{(s)}$ (2%) | $2.5^{\dagger}$ |
| $d_{\rm CH}$ (Å) | $9.44 \pm 0.16\,(1.7\%)$ | 9.4 (2%) | n.a. |
| $\delta_{\rm CH}$ (Å) | 2.8† | 2.8† | n.a. |
| $\delta_{\rm CH3}$ (Å) | $2.84 \pm 0.13\,(5\%)$ | 2.79 (2%) | 3.00 (5%) |
| $r_{\rm P}$ | $0.142 \pm 0.006\,(4\%)$ | $0.133^{(s)}$ (2%) | $0.14^{\dagger}$ |
| $r_{\rm CG}$ | $0.57 \pm 0.02\,(4\%)$ | $0.51^{(s)}$ (2%) | $0.51^{\dagger}$ |
| $r_{12}$ | $0.80 \pm 0.03\,(4\%)$ | $0.76^{(s)}$ (2%) | $0.80^{\dagger}$ |
| $r_{32}$ | $1.98 \pm 0.05\,(3\%)$ | $2.00^{(s)}$ (2%) | $2.09^{\dagger}$ |
| $T$ (°C) | 50† | - | - |
| $V_{\rm H}$ (Å$^3$) | $247 \pm 5\,(1.8\%)$ | $245^{\dagger}$ | $249.6^{\dagger}$ |
| $V_{\rm BW}$( Å$^3$) | $30.24 \pm 0.04\,(0.14\%)$ | n.a. | 29.6 (6%) |
| **Calculated parameters** | | | |
| $z_{\rm ENX}$ (Å) | $23 \pm 3^{\ddagger}$ (15%) | 21.8 (2%) | 23.8 (3%) |
| $z_{\rm P}$ (Å) | $19.3 \pm 0.3^{\ddagger}$ (1.4%) | 20.0 (2%) | 18.8 (8%) |
| $z_{\rm CG}$ (Å) | $16.2 \pm 0.2^{\ddagger}$ (1.2%) | 15.7 (2%) | 17.0 (8%) |
| $V_{\rm L}$ (Å$^3$) | $1189.45^{\ddagger}$ | $1189.40^{\dagger}$ | $1193.80^{\dagger}$ |
| $a_L$ (Å$^2$) | $61.8 \pm 0.9^{\ddagger}$ (1.4%) | 61.3, (2%) | 60.5 (2%) |
| $D_B$ (Å) | $38.1 \pm 0.5^{\ddagger}$ (1.3%) | $38.8^{\ddagger}$ (2%) | $39.5^{\ddagger}$ (3%) |
| $D_{\rm pp}$ (Å) | $36.4^{\ddagger}$ (2%) | $38.7^{\ddagger}$ (2%) | $36.0^{\ddagger}$ (3%) |
| $D_{\rm H1}$ (Å) | $4.1 \pm 0.2^{\ddagger}$ (5%) | $3.96^{\ddagger}$ (2%) | $2.4^{\ddagger}$ (20%) |
| $n_w$ | $\sim 12^{\ddagger}$ | n.a. | $14^{\ddagger}$ (10%) |

$^{\dagger}$ fixed values; $^{\ddagger}$ calculated values; $^{(s)}$ 'soft' Bayesian constrains (Kučerka et al., 2015);

Table S4: List of positional arguments in the `LUV_DMPC` model used as adjustable or fixed parameters ($y$) along with the associate Gaussian or non-informative priors (i.e., hand boundaries $[min, max]$).

| **Model parameters** | | **Prior initialization** | | | **Ref./Exp.** |
|---|---|---|---|---|---|
| $y$ | Description | $\mu_y$ | $\varsigma_y/\mu_y$ | $[min, max]$ | |
| $\mathcal{N}$ | scaling factor | $10^{-7\dagger}$ | - | - | - |
| $n_{\rm v}$ | LUV concentration (number density Å$^{-3}$) | - | - | $[0.9, 3.5]$ | - |
| $R_{\rm v}$ | LUV radius (Å) | 450 | 0.10 | - | $[480, 610]$ |
| $Z$ | LUV polydispersity-related term | - | - | $[10, 20]$ | - |
| $d_{\rm cholCH3}$ | Relative position of group cholCH3 (Å) | 1.0 | 0.03 | - | (Kučerka et al., 2011)[a] |
| $\delta_{\rm cholCH3}$ | Fluctuation width of group cholCH3 (Å) | 2.98 | 0.02 | - | (Kučerka et al., 2011)[a] |
| $d_{\rm PCN}$ | Relative position of group PCN (Å) | 4.0 | 0.08 | - | (Kučerka et al., 2011)[a] |
| $\delta_{\rm PCN}$ | Fluctuation width of group PCN (Å) | 2.77 | 0.20 | - | (Kučerka et al., 2011)[a] |
| $d_{\rm CG}$ | Relative position of group CG (Å) | 0.8 | 0.08 | - | (Kučerka et al., 2011)[a] |
| $\delta_{\rm CG}$ | Fluctuation width of group CG (Å) | 2.43 | 0.20 | - | (Kučerka et al., 2011)[a] |
| $2D_{\rm C}$ | Hydrophobic membrane thickness (Å) | 24.8 | 0.02 | - | (Kučerka et al., 2011) |
| $\sigma_{\rm C}/2D_{\rm C}$ | relative membrane thickness fluctuation | - | - | $[0.01, 0.06]$ | - |
| $\delta_{\rm CH2}$ | Fluctuation width of group $CH_2$ (Å) | 2.45 | 0.05 | - | (Kučerka et al., 2011)[a] |
| $\delta_{\rm CH3}$ | Fluctuation width of group $CH_3$ (Å) | 1.93 | 0.02 | - | (Kučerka et al., 2011) |
| $r_{\rm PCN}$ | PCN relative head-group volume | 0.290 | 0.05 | - | (Kučerka et al., 2011) |
| $r_{\rm CG}$ | CG relative head-group volume | 0.430 | 0.05 | - | (Kučerka et al., 2011) |
| $r_{32}$ | $CH_3$ to $CH_2$ volume ratio | 1.93 | 0.05 | - | (Kučerka et al., 2011) |
| $T$ | Sample temperature (°C) | $50^\dagger$ | - | - | - |
| $V_{\rm H}$ | Lipid head-group volume (Å$^3$) | 324 | 0.02 | - | (Nagle et al., 2019)[b] |
| $V_{\rm BW}$ | Hydration-layer water volume (Å$^3$) | - | - | $[28.5, 31.5]$ | - |
| $Const.$ | Constant background | - | - | $[1.7 \times 10^{-3}, 2.8 \times 10^{-3}]$ | - |

[a] Priors width are set to be consistent with the choices for POPC and POPE systems;
[b] Derived from the range of values reported for PC lipids;
† Fixed parameter.

Table S5: Results of the analysis of DMPC at 50 °C with the LUV_DMPC model. Best values and relative uncertainties are reported as median ± MAD (Median Absolute Deviation) and compared with previous results. Number of iterations $N_I = 300$. Note that relative uncertainties in previous works refer to standard deviations.

| **Parameter** $y$ | median±MAD | Ref. (Kučerka et al., 2011) |
|---|---|---|
| $R_{\rm v}$ (Å) | $552 \pm 6$ (1.1%) | n.a. |
| $Z$ | $24.1 \pm 1.6$ (7%) | n.a. |
| $d_{\rm CholCH3}$ (Å) | $1.00 \pm 0.03$ (3%) | $1.0^‡$ |
| $\delta_{\rm CholCH3}$ (Å) | $2.98 \pm 0.06$ (1.9%) | $2.98^†$ |
| $d_{\rm PCN}$ (Å) | $3.9 \pm 0.3$ (7%) | $4.0^‡$ |
| $\delta_{\rm PCN}$ (Å) | $3.05 \pm 0.08$ (3%) | 2.77 (2%) |
| $d_{\rm CG}$ (Å) | $0.82 \pm 0.06$ (7%) | $0.8^‡$ |
| $\delta_{\rm CG}$ (Å) | $2.20 \pm 0.18$ (8%) | 2.43 (2%) |
| $2D_{\rm C}$ (Å) | $24.6 \pm 0.3$ (1.4%) | $24.8^‡$ (2%) |
| $\varepsilon_{\rm C}$ | $0.027 \pm 0.002$ (8%) | n.a. |
| $\delta_{\rm CH2}$ (Å) | $2.48 \pm 0.11$ (4%) | $2.45^{(s)}$ (2%) |
| $\delta_{\rm CH3}$ (Å) | $1.93 \pm 0.03$ (1.8%) | 1.93 (2%) |
| $r_{\rm PCN}$ | $0.291 \pm 0.013$ (4%) | $0.290^{(s)}$ (2%) |
| $r_{\rm CG}$ | $0.4143 \pm 0.014$ (3%) | $0.43^{(s)}$ (2%) |
| $r_{32}$ | $1.91 \pm 0.06$ (3%) | $1.93^{(s)}$ (2%) |
| $T$ (°C) | 50† | - |
| $V_{\rm H}$ (Å$^3$) | $321 \pm 5$ (2%) | $331^†$ |
| $V_{\rm BW}$( Å$^3$) | $30.45 \pm 0.12$ (0.4%) | n.a. |
| **Calculated parameters** | | |
| $z_{\rm CholCH3}$ (Å) | $18.0 \pm 1.6^‡$ (6%) | 18.2 (2%) |
| $z_{\rm PCN}$ (Å) | $17.0 \pm 0.3^‡$ (2%) | 17.2 (2%) |
| $z_{\rm CG}$ (Å) | $13.10 \pm 0.12^‡$ (0.9%) | 13.2 (2%) |
| $V_{\rm L}$ (Å$^3$) | $1115.2^‡$ | $1115.0^†$ |
| $a_L$ (Å$^2$) | $64.6 \pm 0.9^‡$ (1.4%) | 63.3, (2%) |
| $D_B$ (Å) | $34.5 \pm 0.5^‡$ (1.4%) | $35.2^‡$ (2%) |
| $D_{\rm pp}$ (Å) | $29.9^‡$ (2%) | $32.2^‡$ (2%) |
| $D_{\rm H1}$ (Å) | $6.6 \pm 0.2^‡$ (3%) | $3.73^‡$ (2%) |
| $n_w$ | $\sim 17^‡$ | n.a. |

† fixed values; ‡ calculated values;
$^{(s)}$ 'soft' Bayesian constrains (Kučerka et al., 2011).

#### S3 Program output

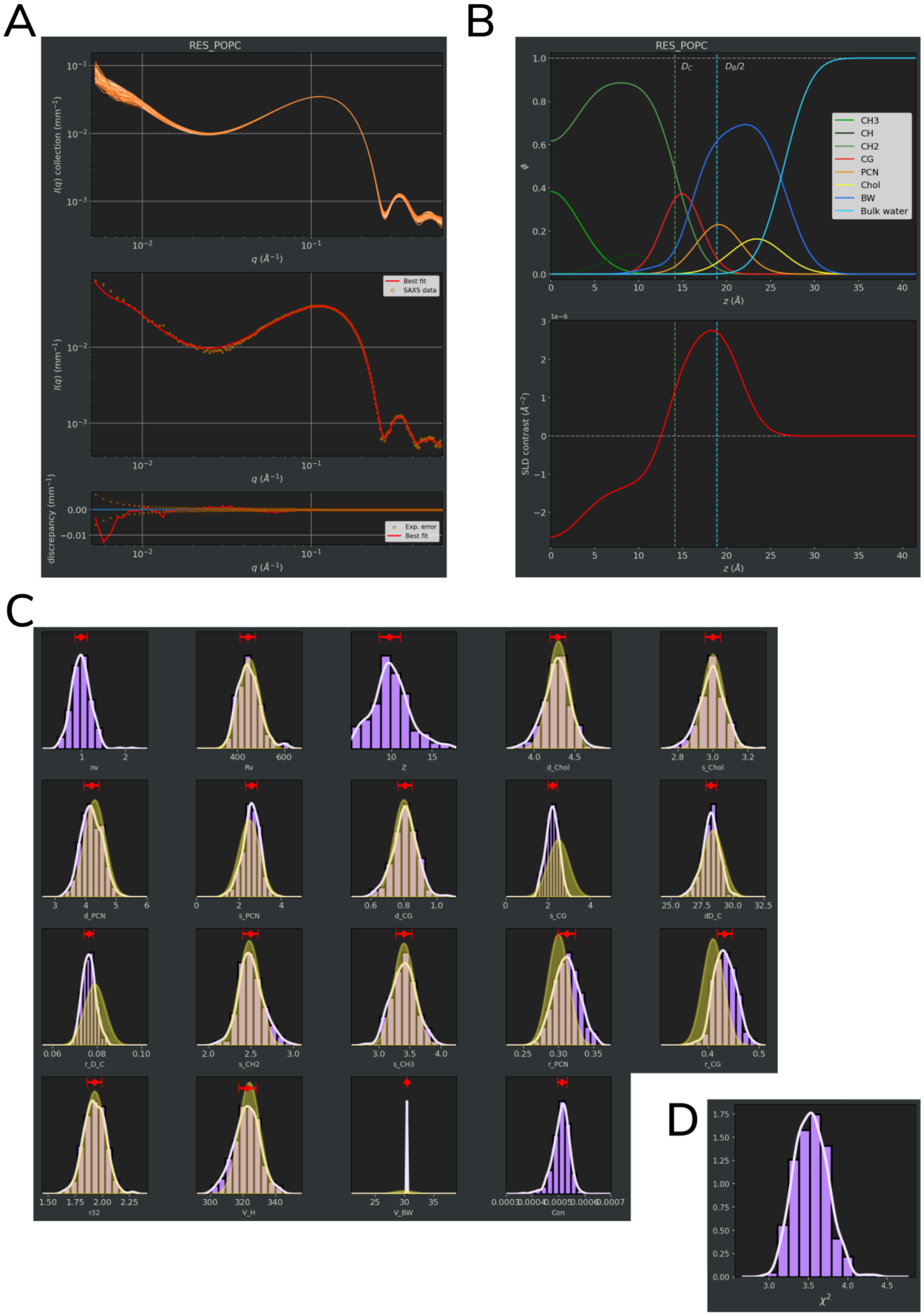


Figure S2: *SAS_MoCa* output image files for the analysis of POPC systems. *plot.png* (A), *plot_SDP-SLD.png* (B), *plot_histograms.png* (C) and *plot_histogram_X2.png* (D).

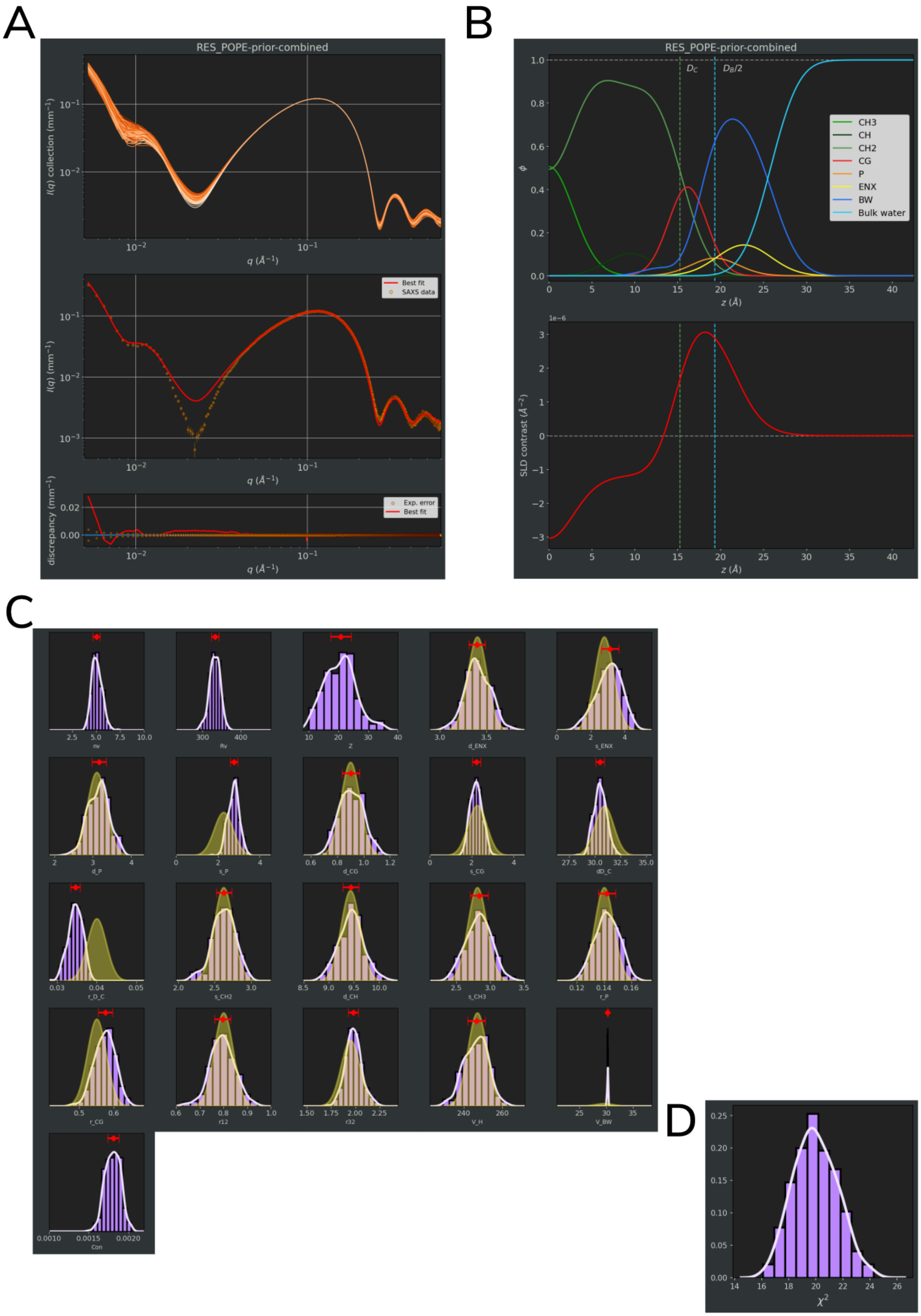


Figure S3: *SAS_MoCa* output image files for the analysis of POPE systems. *plot.png* (A), *plot_SDP-SLD.png* (B), *plot_histograms.png* (C) and *plot_histogram_X2.png* (D).

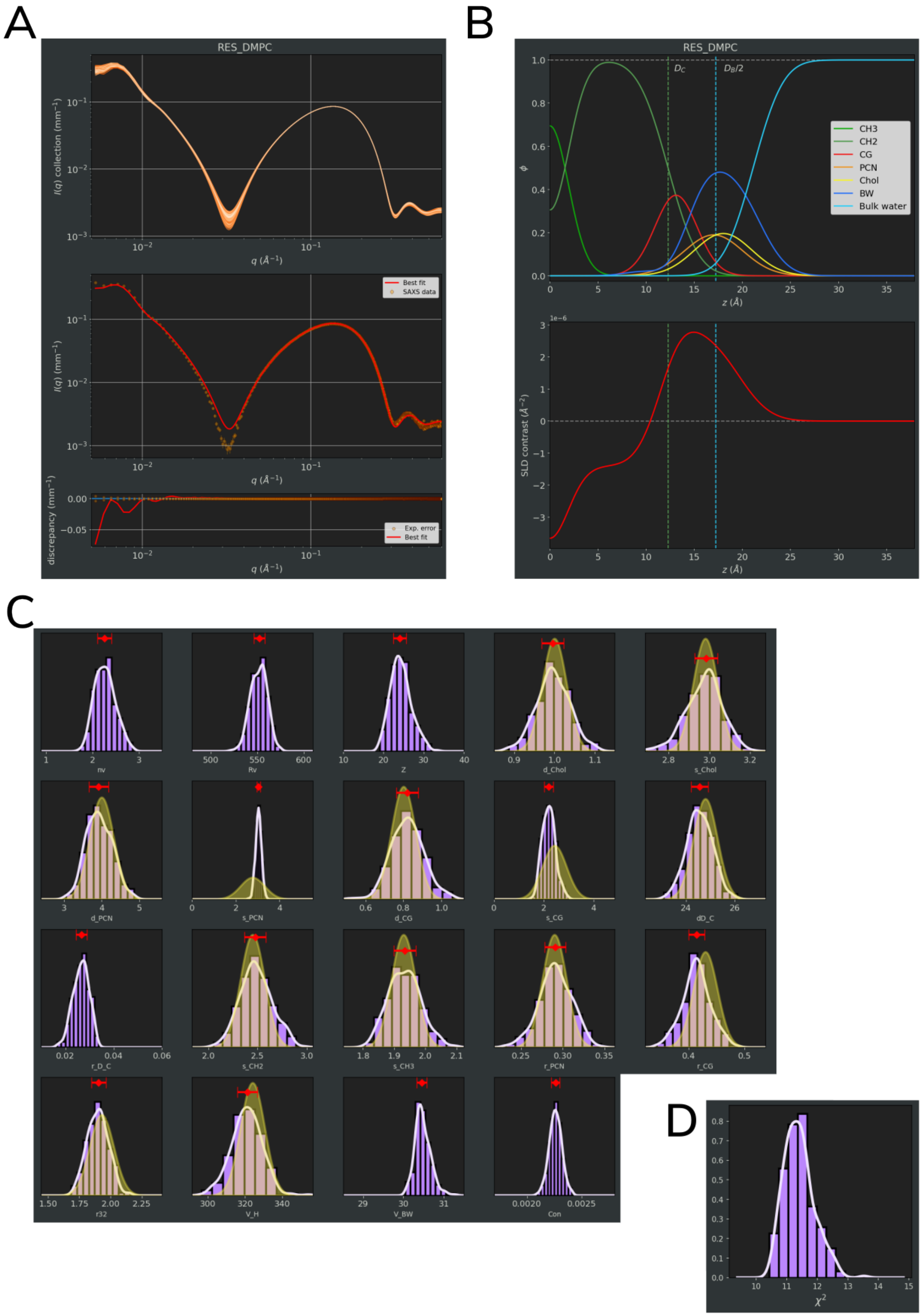


Figure S4: *SAS_MoCa* output image files for the analysis of DMPC systems. *plot.png* (A), *plot_SDP-SLD.png* (B), *plot_histograms.png* (C) and *plot_histogram_X2.png* (D).

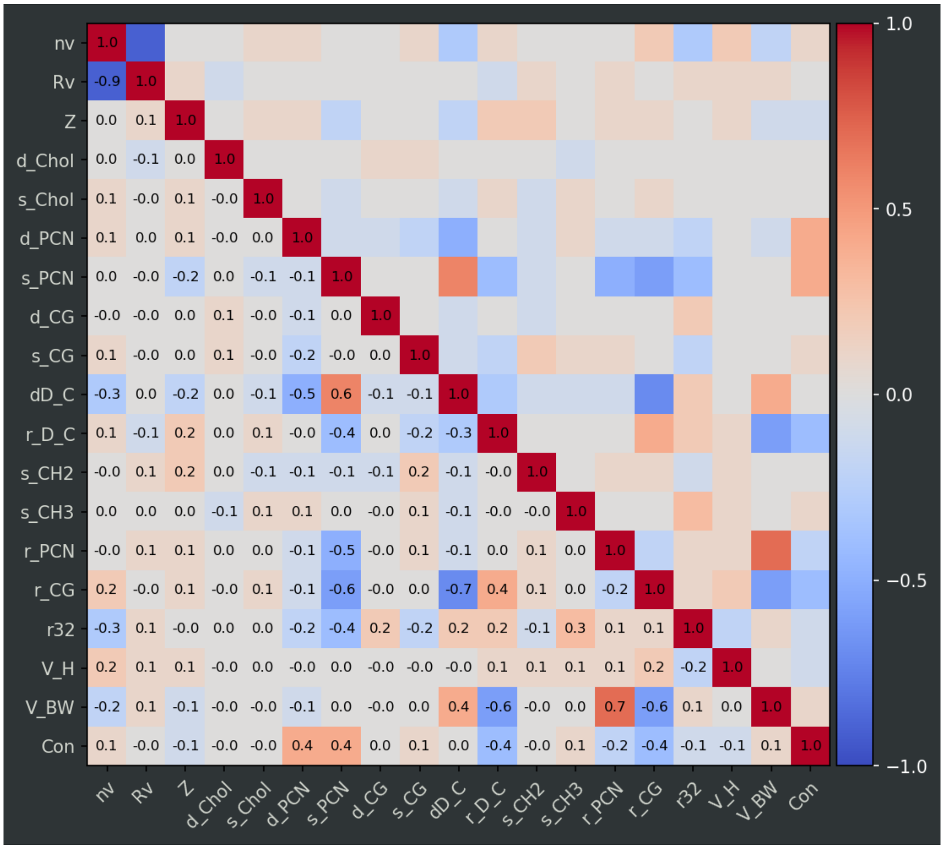


Figure S5: *SAS_MoCa* output image file *plot_correlations.png* for the analysis of POPC systems.

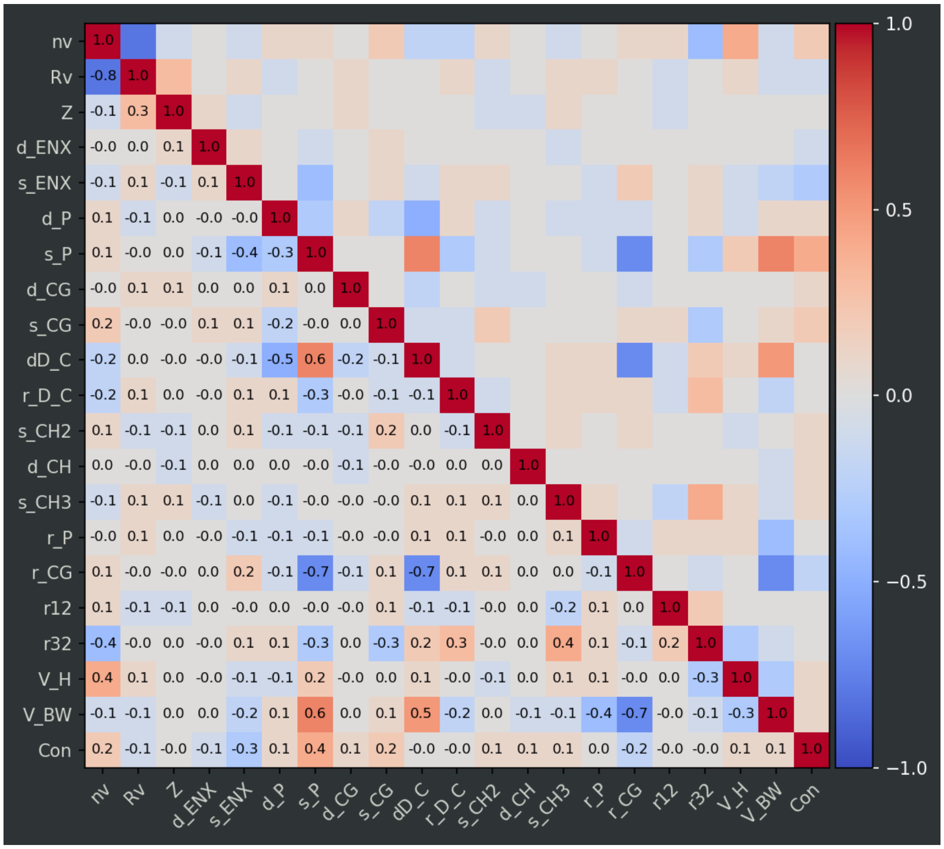


Figure S6: *SAS_MoCa* output image file *plot_correlations.png* for the analysis of POPE systems.

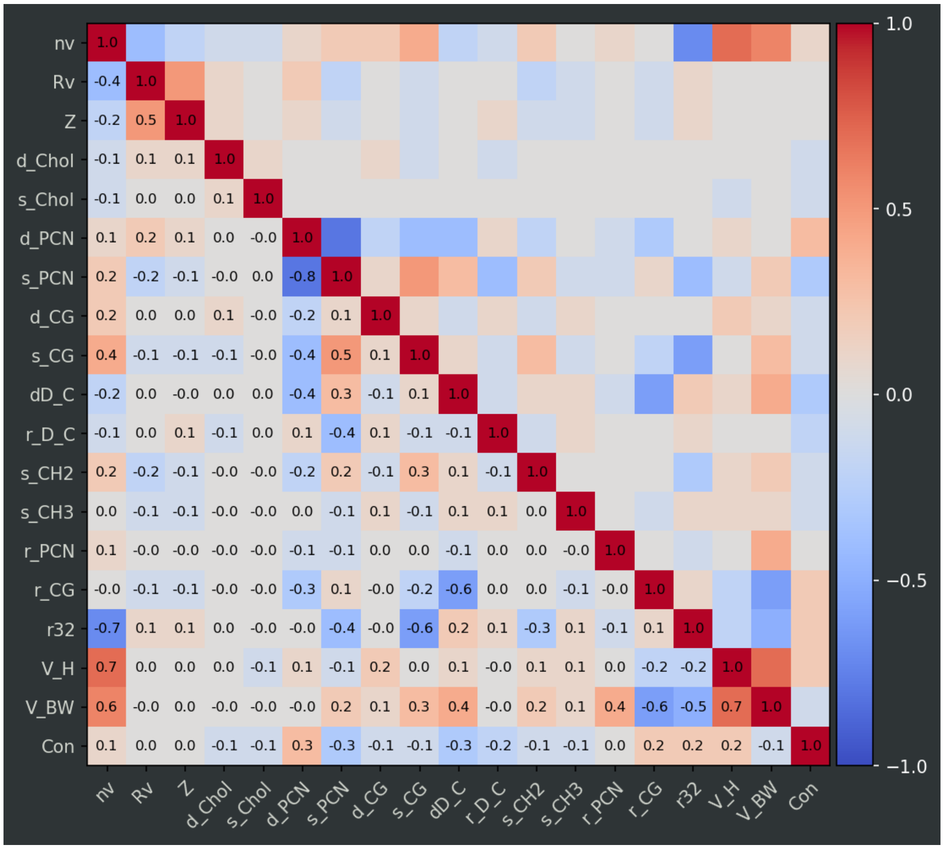


Figure S7: *SAS_MoCa* output image file *plot_correlations.png* for the analysis of DMPC systems.
